# Complex Intracellular Mechanisms of TBK1 Kinase Activation Revealed by a Specific Small Molecule Inhibitor

**DOI:** 10.1101/2022.10.11.511671

**Authors:** Junqiang Ye, Noah Dephoure, Tom Maniatis

## Abstract

TANK-binding kinase 1 (TBK1), a dimeric serine/threonine protein kinase, plays a critical role in multiple signaling pathways including innate immunity, autophagy and cell death. TBK1 is activated by the phosphorylation of an essential serine residue 172 (S172) within the activation loop of the kinase domain, and this phosphorylation can occur by trans-autophosphorylation: one TBK1 dimer phosphorylates a second dimer at S172. Here we show that phosphorylation of TBK1 S172 in cultured human cells in response to multiple inducers is reduced, but not abolished by the highly specific and potent TBK1 small molecule inhibitor GSK8612. Thus, upstream kinase(s) must phosphorylate TBK1 in response to inducers in cultured cells. We show that distinct upstream kinases are recruited for the activation of TBK1 in response to different inducers. We also identify extensive crosstalk among TBK1, IKKβ and IKKε kinases in the cellular response to various inducers. In addition, we show that dsDNA and dsRNA trigger dynamic intracellular translocation of TBK1, leading to its localization and activation on the Golgi apparatus or mitochondria, respectively. GSK8612 does not block the intracellular localization of TBK1. We conclude that TBK1 is activated by both upstream kinase phosphorylation and by trans-autophosphorylation, and the signal-dependent spatial engagement of TBK1 with other signaling molecules is a fundamental mechanism for its specific activation by multiple inducers.

## Introduction

The serine/threonine protein kinase TANK-binding kinase 1 (TBK1) plays critical roles in cellular responses to microbial infection, inflammation and autophagy (1, 2). TBK1 is also required for cell cycle and cell death regulation, and has been implicated in oncogenesis (3). TBK1 is remarkably pleiotropic, as mutations of TBK1 cause or associate with a number of human diseases, including amyotrophic lateral sclerosis (ALS) and frontotemporal dementia (FTD), normal-tension glaucoma (NTG), systemic autoinflammation, childhood herpes simplex virus-1 encephalitis (HSE), and more recently severe COVID-19 disease progression (4-9).

TBK1 is present as a protein dimer of identical subunits in vivo and is inactive in the resting state. However, in response to various activating signals, TBK1 is phosphorylated at serine residue 172 (S172) located within the activation loop of the kinase domain (KD), a key step required to trigger and lock the KD in an active conformation (10-13). Phospho-S172 coordinates with arginine residues within the activation loop (R162), catalytic loop (R134) and C-helix (R54), leading to the repositioning of the C-helix and reorganization of the activation loop from the inactive conformation. Thus, the kinase is enabled for substrate binding and the subsequent phosphoryl transfer step (11). Phosphorylation of multiple distinct target proteins by TBK1 leads to distinct and specific biological outcomes.

A remarkable feature of TBK1 is its substrate specificity in response to different upstream signals (3). For example, TBK1 activated by virus infection specifically phosphorylates the transcription factor IRF3 (critical for the innate immune response), which leads to IRF3 nuclear translocation and expression of the interferon-β gene (14, 15). DNA and RNA viruses activate TBK1 through distinct pathways, with the former signaling through cGAS-STING pathway, and the latter through the RIG-I/MDA5-MAVS pathway (16). However, TBK1 activated by either DNA or RNA viruses phosphorylates IRF3. By contrast, TBK1 activated during selective autophagy phosphorylates the autophagy receptors SQSTM1/p62 and OPTN (1), while TBK1 activated by TNFα signaling phosphorylates RIPK1 (17, 18). TBK1 kinase activity is restricted to distinct substrate proteins within specific pathways. For example, TBK1 activated during mitophagy (selective degradation of damaged mitochondria through autophagy) does not act on IRF3 or RIPK1(19). Moreover, TNFα activated TBK1 does not phosphorylate IRF3 (20). Similarly, viral infection does not necessarily lead to autophagy receptor phosphorylation when the autophagy pathway is not activated (for example, certain RNA viruses, such as Sendai virus, are poor inducers of autophagy). The basis of this remarkable TBK1 specificity in distinct signaling pathways is not fully understood. Chen and colleagues showed that adaptor protein phosphorylation confers selectivity in innate immunity (20). Specifically, conserved serine/threonine sites within STING in the DNA pathway, MAVS in the RNA pathway and TRIF in TLR3/4 signaling are phosphorylated by TBK1 and IKK kinases upon activation of their respective pathways. Phosphorylated sites on these adaptor proteins function as docking sites for the recruitment and phosphorylation of IRF3. It is important to note that STING is only phosphorylated during double-stranded DNA (dsDNA) signaling, not during double-stranded RNA (dsRNA) signaling, despite the fact that TBK1 is activated in both pathways (20). TBK1 appears to be recruited to distinct signaling complexes in response to specific inducers, and phosphorylates target proteins within the complex (3), however, the molecular mechanisms of specific recruitment of TBK1 to different pathways are not known.

A significant gap in our understanding of these signaling events is the mechanism of TBK1 activation. As mentioned above, TBK1 is activated by the phosphorylation of S172 residue within its activation loop. Biochemical and structural studies utilizing over-expression experiments and recombinant proteins have led to a trans-autophosphorylation model of TBK1 activation (10, 11) based on two key observations: 1) TBK1 can efficiently phosphorylate other TBK1 molecules in kinase assays and over-expression experiments. One caveat in these experiments, however, is the presence of S172 phosphorylated TBK1 molecules prior to induction (10, 11); and 2) TBK1 subunits within the same dimer cannot phosphorylate each other, as they face away from each other (based on the TBK1 crystal structure) (11-13). The trans-autophosphorylation model was proposed to account for many signaling pathways involving TBK1. In all these cases, the initial TBK1 S172 phosphorylation step, which is key to turning on TBK1 kinase activity, remains elusive. A prevalent view is that the initial S172 phosphorylation is carried out by other TBK1 molecules recruited to the same signaling complex (17, 19, 20). The difficulty with this model is that the conformations of active and inactive TBK1 are different, particularly regarding the C-helix and the activation loop within the kinase domain (10, 11). Thus, it is not clear how an inactive TBK1 dimer becomes active without S172 phosphorylation, which, based on structural studies, is required for the kinase to adopt the active confirmation for substrate binding (11). If this does occur, it is probably very inefficient (10). An alternative, and more likely hypothesis is that the initial TBK1 S172 phosphorylation step is carried out by an upstream kinase(s). Previous studies suggest that TBK1 can be phosphorylated by other kinases (19, 21, 22), but the nature and mechanism of these activities have not been carefully investigated.

To understand the mechanisms of TBK1 activation and its signal dependent recruitment, we carried out experiments using the highly specific TBK1 small molecule inhibitor GSK8612 (23), and challenged cultured cells with dsDNA or dsRNA, two related but distinct inducers of TBK1 activation. Combining chemical biology, genetically modified cell lines, cellular imaging and biochemistry, we found that there are distinct upstream kinases that phosphorylate TBK1 in response to dsDNA and dsRNA induction. TBK1 intracellular localization is dynamically regulated upon stimulation, with dsDNA induced TBK1 enrichment on Golgi, and dsRNA induced enrichment on mitochondria, respectively. Distinct TBK1 inducers, such as dsDNA or dsRNA, can lead to TBK1 activation at distinct subcellular locations. For example, TBK1 is engaged in different complexes with STING or p62 after dsDNA challenge.

We also show that IKKε, a kinase closely related to TBK1, is recruited to TBK1-STING signaling complexes upon dsDNA stimulation, and partially contributes to TBK1 phosphorylation. In addition, we find that the IKKβ kinase is recruited to the TBK1 signaling complex upon dsRNA stimulation, and contributes to TBK1 and IRF3 phosphorylation. Finally, we identify a small molecule 7DG as a TBK1 modulator, which induces a striking perinuclear localization of TBK1. 7DG in combination with GSK8612 strongly inhibits dsDNA-induced TBK1 activation, demonstrating that the full activation of TBK1 requires a 7DG sensitive kinase and trans-autophosphorylation. Taken together, these studies reveal a remarkably complex network of signaling pathways in which a single kinase can play a central role.

## Results

### Upstream kinases are involved in the activation of TBK1 in multiple signaling pathways

Based on the central role of TBK1 in critical signaling pathways, several small molecule inhibitors of TBK1, including BX795, MRT67307, and more recently GSK8612, have been identified, developed and applied (21-23). These small molecule inhibitors have a common 2,4-diaminopyrimidine core attached to different chemical groups, which determine binding specificity. BX795 and MRT67307 have been widely used, based on their early availability, to identify specific signaling pathways that require TBK1 kinase activity.

We pretreated cultured human epithelial HT1080 cells with a 10µM concentration of each inhibitor, a concentration known to strongly inhibit TBK1 kinase activity (21-23). We then carried out inductions by transfecting cells with dsDNA (poly dG:dC or dI:dC) or dsRNA (poly I:C) by lipofectamine. Cells were harvested after 2 hrs and the phosphorylation of various proteins determined by Western blots probed with phospho-specific antibodies. Both dsDNA and dsRNA robustly induce TBK1 phosphorylation at S172 (pTBK1) in cells that were not treated with any inhibitors (Fig. 1A, dsDNA appears to be a stronger inducer of TBK1 activation compared to dsRNA in this cell line). TBK1 is known to phosphorylate several target proteins under these conditions (24, 25), including p62, which is phosphorylated at S403 by TBK1 upon either dsDNA or dsRNA stimulation. As expected, all of the small molecule inhibitors abolished p62 phosphorylation (Fig. 1A), demonstrating p62 S403 phosphorylation is carried out exclusively by TBK1 under these conditions. By contrast, TBK1 S172 phosphorylation was not completely inhibited in cells pretreated with inhibitors followed by dsDNA/dsRNA stimulation (Fig. 1A, top two blots). For example, in cells treated with GSK8612, an approximately 50% reduction of pTBK1 was observed compared to control cells after dsDNA treatment, and less of a reduction was observed after dsRNA treatment (Fig. 1A), revealing different levels of autophosphorylation of TBK1 in response to different inducers. By comparison, treatment with BX795 or MRT67307 led to a higher level of pTBK1 in stimulated samples compared to controls. Unexpectedly, treating cells with these two inhibitors alone, in the absence of any other stimulator, induced pTBK1, suggesting that although the small molecules inhibited TBK1, they apparently activated other kinases which phosphorylate TBK1 at S172. Additional experiments revealed that 0.5-2µM of MRT67307 alone was sufficient to induce pTBK1 in HT1080 and HeLa cells (Supplementary Fig.1A, 1B). In contrast, GSK8612 alone did not induce TBK1 phosphorylation. This compound abolished the low basal level of pTBK1 in control cells (Fig. 1A). We also observed that both BX795 and MRT67307 increased the expression of A20 protein, and induced cleavage of OPTN (even without inducers), providing additional evidence of off-target effects of these small molecules. Finally, we found that BX795 blocked the dsRNA-induced activation of JNK (Fig. 1A), consistent with previously reported off-target results (21).

**Figure 1.**
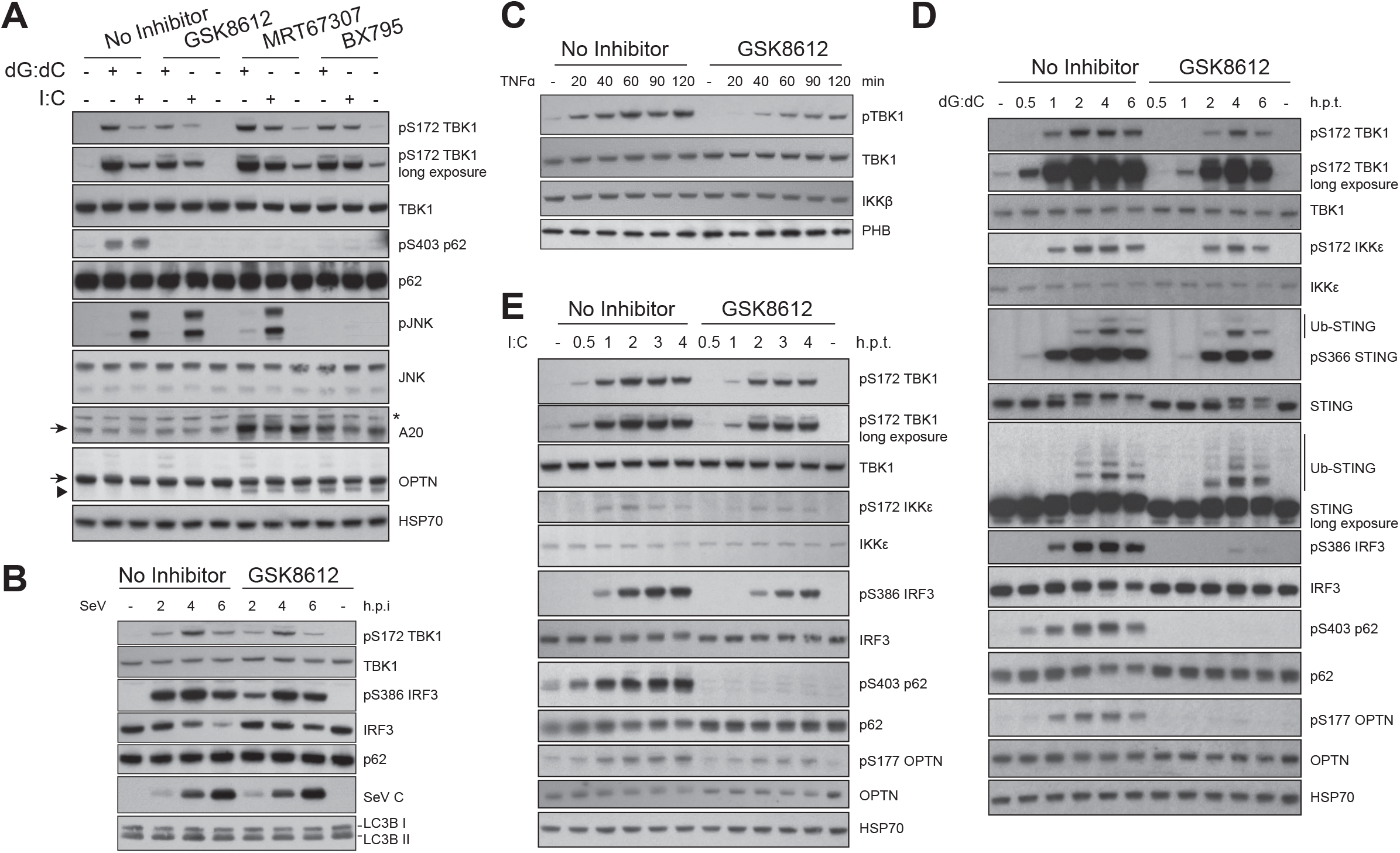
Upstream kinases are involved in the activation of TBK1. **A**. GSK8612 is a more specific TBK1 inhibitor compared to BX795 and MRT67307. HT1080 cells were either untreated, or pretreated with GSK8612, MRT67307 or BX795 (10µM) for 1hr, then treated by lipofectamine 2000 mediated transfection of dsDNA (poly dG:dC, 1.6µg/ml) or dsRNA (poly I:C, 1.6µg/ml) for 2hrs. Total cell lysates were prepared, and the phosphorylation and expression of various proteins determined by Western blotting with specific antibodies. A nonspecific band from A20 blot is labeled with an asterisk. The target bands of A20 and OPTN are marked by arrows, and a cleavage product of OPTN is labeled by an arrowhead. **B**. Kinetic study of HT1080 cells infected with Sendai virus in the absence or presence of GSK8612. Control cells or cells pretreated with GSK8612 (10µM) for 1hr were infected with Sendai virus (300 HAU/ml) for increasing time (2, 4 and 6hrs). Total cell lysates were prepared at specified time points post infection, and the phosphorylation and expression of TBK1, IRF3, p62, Sendai virus C protein and LC3B were determined by Western blotting with specific antibodies. **C**. Kinetic study of HT1080 cells stimulated with TNFꭤ in the absence or presence of GSK8612. Control cells or cells pretreated with GSK8612 (10µM) for 1hr were treated with TNFꭤ (10ng/ml) for increasing time (20 to 120 minutes). Total cell lysates were prepared at specified time points post stimulation, and the profile of phosphorylation and expression of various proteins were determined by Western blotting. **D**. Kinetic study of HT1080 cells stimulated with dsDNA in the absence or presence of GSK8612. Control cells or cells pretreated with GSK8612 (10µM) for 1hr were challenged with dsDNA (poly dG:dC) for increasing time (30 minutes to 6hrs). Total cell lysates were prepared at specified time post stimulation. The phosphorylation profile and expression of various proteins were determined by Western blotting with specific antibodies. **E**. Kinetic study of HT1080 cells stimulated with dsRNA in the absence or presence of GSK8612. Control cells or cells pretreated with GSK8612 (10µM) for 1hr were challenged with dsRNA (poly I:C) for increasing time (30 min to 4hrs). Total cell lysates were prepared at specified time points post stimulation. The expression and phosphorylation of various proteins were determined by Western blotting with specific antibodies.

These experiments show that GSK8612 is the most specific TBK1 inhibitor currently available. Given these results, previous studies implicating TBK1 kinase activity of specific substrates using BX795 or MRT67307 must be considered in the light of their lack of specificity. Most importantly, however, these data unequivocally show that a yet to be identified kinase(s) upstream of TBK1 phosphorylates TBK1 in response to dsDNA and dsRNA.

As a further test of upstream TBK1 kinases, we monitored TBK1 S172 phosphorylation after Sendai virus (SeV) infection (Fig. 1B), or TNFα stimulation (Fig. 1C). Both SeV and TNFα induced TBK1 S172 phosphorylation in the presence of GSK8612. In the case of SeV infection, pTBK1 levels were only slightly reduced by GSK8612 compared to the equivalent time points in infected control cells (Fig. 1B), while this inhibitor suppressed greater than 50% of pTBK1 after TNFα stimulation (Fig. 1C). These observations clearly show that the proposed TBK1 trans-autophosphorylation mechanism alone is insufficient to explain the full activation of TBK1 in human cells in culture.

### Signal-dependent engagement of TBK1 with its target proteins

We carried out kinetic studies of cells treated with either dsDNA or dsRNA in the presence or absence of GSK8612, and monitored the phosphorylation of p62, IRF3, and OPTN, all well-established TBK1 target proteins, at different times post stimulation. Like our earlier observations, S403 phosphorylation of p62 induced by dsDNA or dsRNA was completely inhibited by GSK8612 treatment (Fig. 1D, 1E), demonstrating the potency of this compound. We also found that IRF3 phosphorylation was strongly inhibited by GSK8612 treatment in dsDNA induced samples (Fig. 1D). Surprisingly, however, IRF3 phosphorylation was still observed in dsRNA treated samples, but was delayed and reduced, with a level of less than 50% of that in control cells at equivalent time points (Fig 1E). Similarly, SeV-induced IRF3 phosphorylation was reduced, but not blocked by GSK8612 pretreatment (Fig. 1B). Moreover, dsDNA-induced OPTN S177 phosphorylation was also inhibited by GSK8612 but was delayed and reduced (<50% of control level) in dsRNA treated samples as well (Fig. 1D, 1E). Since TBK1 kinase activity was strongly inhibited by GSK8612, a different kinase(s) must phosphorylate IRF3 and OPTN when cells were treated with dsRNA in the presence of GSK8612.

TBK1 is known to contribute to STING S366 phosphorylation (20, 26), which is specifically induced in cells stimulated with dsDNA (27). In GSK8612 treated samples, STING S366 phosphorylation was delayed and reduced (from far less than 50% at early time points to about 50% the level in control cells at later time points), but not abolished (Fig. 1D). Interestingly, the multiple bands of STING on SDS-PAGE, presumably due to ubiquitination, were not affected by GSK8612 treatment. The turnover of STING also appeared to be unaffected by GSK8612 treatment (Fig. 1D).

These data show that TBK1 engages its target proteins in a signal-dependent manner: dsDNA-induced IRF3 and OPTN phosphorylation is primarily carried out by TBK1. By contrast, dsRNA and SeV can activate other kinases in addition to TBK1, which can phosphorylate IRF3 and OPTN in the absence of catalytically active TBK1 (inhibited by GSK8612). The phosphorylation of p62 at S403, however, occurs exclusively through TBK1 following either dsDNA or dsRNA induction. The same observation was made in HeLa cells (Supplementary Fig. 1C). These results suggest that TBK1 activated by different stimuli is recruited to distinct signaling complexes, which are comprised of unique or partially overlapping protein components under different conditions. Remarkably, inhibiting TBK1 in these distinct complexes can lead to different outcomes, depending on the presence of other, yet-to-be identified kinases that can act on targets previously thought-to-be phosphorylated only by TBK1.

### TBK1 dynamics in dsDNA and dsRNA signaling

The biochemical data presented here show that TBK1 recruitment to signaling complexes and engagement with target proteins as signal-dependent. To study the spatial distribution/dynamics of TBK1 in response to different inducers, we carried out immunofluorescence (IF) staining of cells challenged with different inducers, using antibodies specific for total TBK1 or pTBK1, in conjunction with antibodies for selected subcellular markers.

We first asked whether inducers change the intracellular distribution of TBK1 using an antibody that recognizes total TBK1 protein (binding to the C-terminus of TBK1, the specificity of this antibody was validated in Supplementary Fig. 2). We found that dsDNA stimulation of HT1080 cells for 2hrs led to TBK1 aggregation in the cytosol, which co-stained with the Golgi marker GM130 (Fig. 2A). In contrast, TBK1 was relatively evenly distributed in the cytosol under resting conditions, or after being treated by GSK8612, demonstrating that TBK1 localization to the Golgi was dsDNA-stimulation specific (Fig. 2A). Staining cells with antibodies for pTBK1 revealed that the majority of pTBK1 also co-localizes with GM130 (Fig. 2A). We noticed that pTBK1 is unevenly distributed in the cytoplasm, with a major fraction concentrated on Golgi, and the remaining in a punctate pattern dispersed in the cytosol (Fig. 2A). The non-uniform spatial distribution of pTBK1 suggests that these pTBK1 signaling complexes are heterogeneous, likely containing different protein components. Consistent with biochemical results, treating cells with dsDNA in the presence of GSK8612 reduced the level of pTBK1. The strong Golgi-associated pTBK1 signal induced by dsDNA in the absence of GSK8612 was not observed after inhibitor treatment, and instead, a more punctate staining pattern of pTBK1 was observed on Golgi (Fig. 2A). Importantly, total TBK1 staining revealed that GSK8612 did not prevent the localization of TBK1 on Golgi, as a similar punctate pattern of total TBK1 distribution was observed after dsDNA stimulation, irrespective of GSK8612 treatment (Fig. 2A). These data indicate that TBK1 localization to Golgi occurs prior to its full activation. Despite a reduction, pTBK1 was observed on Golgi in cells pretreated with GSK8612 followed by dsDNA stimulation, indicating that Golgi-associated kinases can phosphorylate TBK1, a step likely required for TBK1 activation. Under normal circumstances (in the absence of inhibitor), activated TBK1 can phosphorylate other TBK1 molecules recruited to Golgi by a trans-autophosphorylation mechanism, or directly engage with downstream target proteins (28).

**Figure 2.**
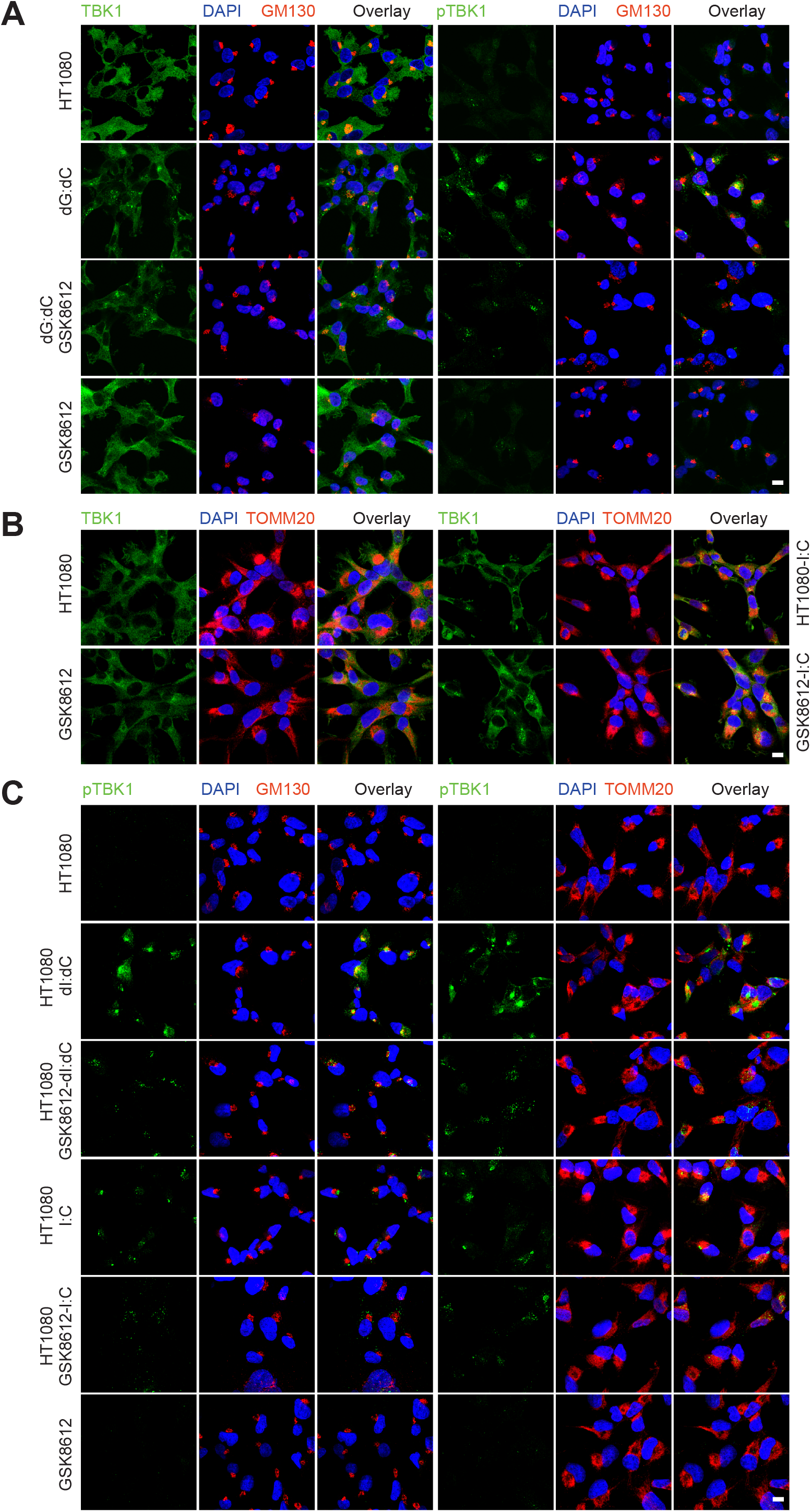
Intracellular TBK1 dynamics upon dsDNA/RNA stimulation. **A**. Phosphorylated TBK1 was enriched on Golgi apparatus upon dsDNA stimulation. HT1080 cells were stimulated with dsDNA (poly dG:dC) for 2hrs in the absence or presence of GSK8612 (10µM, pretreated for 1hr). Cells were fixed and subjected to immunofluorescence staining with antibodies against pTBK1, TBK1 and GM130. Distributions of these proteins were analyzed by confocal microscopy. Scale bar: 10µm. **B**. dsRNA induces mitochondria enrichment of TBK1. HT1080 cells were stimulated with dsRNA (poly I:C) for 2hrs in the absence or presence of GSK8612 (10µM, pretreated for 1hr). Cells were fixed and subjected to immunofluorescence staining with antibodies against TBK1 and TOMM20. Scale bar: 10µm. **C**. Distribution of pTBK1 after dsDNA or dsRNA treatment. HT1080 cells were either untreated or pretreated with GSK8612 (10µM) for 1hr, followed by stimulation with dsDNA (poly dI:dC) or dsRNA (poly I:C) for 2hrs. Cells were fixed and subjected to IF staining with antibodies against pTBK1, GM130 and TOMM20. Scale bar: 10µm.

dsRNA stimulation leads to a similar pattern of total TBK1 as that observed with dsDNA stimulation, as they form aggregates irrespective of the presence of GSK8612 (Fig. 2B). In contrast to Golgi localization of TBK1 after dsDNA stimulation, dsRNA-induced TBK1 aggregates were enriched on mitochondria (stained by TOMM20), consistent with transfected dsRNA signaling through the MAVS protein on mitochondria (29). The reduction of pTBK1 induced by dsRNA in the presence of GSK8612 was smaller compared to its effect on dsDNA stimulation (Fig. 1A, 1E). GSK8612 did not block TBK1 mitochondria localization induced by dsRNA stimulation (Fig. 2B), similar to its ineffectiveness in blocking TBK1 Golgi localization induced by dsDNA stimulation (Fig. 2A).

We also examined the distribution of pTBK1 with respect to Golgi and mitochondria after dsDNA/RNA stimulations. The dominant Golgi-localization of pTBK1 induced by dsDNA filled the gaps between mitochondria (stained by TOMM20) and nucleus (DAPI staining, Fig. 2C), with a small fraction of pTBK1 co-localizing with TOMM20. The staining was primarily away from the nucleus, and more punctate. This activated TBK1 is likely involved in non-STING related signaling events (to be addressed below). As mentioned above, GSK8612 reduced the intensity and size of pTBK1 staining signals on Golgi and was replaced with a punctate small-dotted pTBK1 pattern on Golgi with little co-staining of mitochondria (Fig. 2C). On the other hand, dsRNA induced more co-staining of pTBK1 with TOMM20, with few examples that the pTBK1 also stained positive for GM130 (Fig. 2C). GSK8612 treatment also reduced the size and intensity of pTBK1 signal, but these pTBK1 still associated with mitochondria and did not co-stain for GM130 (Fig. 2C). These data show that dsDNA and dsRNA induce distinct subcellular localization of TBK1, which likely contributes to the distinct outcomes of these two inducers. Although GSK8612 reduces the total level of pTBK1, it does not block the movement of TBK1 to target destination, nor does it inhibit the initial TBK1 phosphorylation, indicating that TBK1 recruitment to specific subcellular sites and its phosphorylation by upstream kinases are an integral part of its activation.

### Involvement of IKKβ in the phosphorylation of TBK1 and IRF3 in dsRNA signaling

Since GSK8612 alone does not block the phosphorylation of TBK1 in response to either dsDNA or dsRNA stimulation, we sought to identify other kinases involved in the activation of TBK1. We took a double-inhibitor approach: cells were pretreated with GSK8612 in combination with another kinase inhibitor, and then challenged with dsDNA/dsRNA for 2hrs. pTBK1 levels under different conditions were monitored by Western blots. Based on our preliminary kinase inhibitor library screening data, six additional kinase inhibitors were chosen for this study: GSK3β inhibitor TWS119, IKKβ inhibitor TPCA-1, PKR inhibitor C16, AKT inhibitor VIII, and two tyrosine kinase inhibitors FLT3-II and SU11652.

Strikingly, the PKR inhibitor, C16, in combination with GSK8612 significantly inhibited dsDNA-induced TBK1 phosphorylation; concomitantly, STING phosphorylation was also abolished by the combined treatment (Fig. 3A). Interestingly, C16 and GSK8612 only partially reduced TBK1 phosphorylation induced by dsRNA (Fig. 3B), showing that different upstream kinases are involved in the phosphorylation of TBK1 in response to dsDNA and dsRNA.

**Figure 3.**
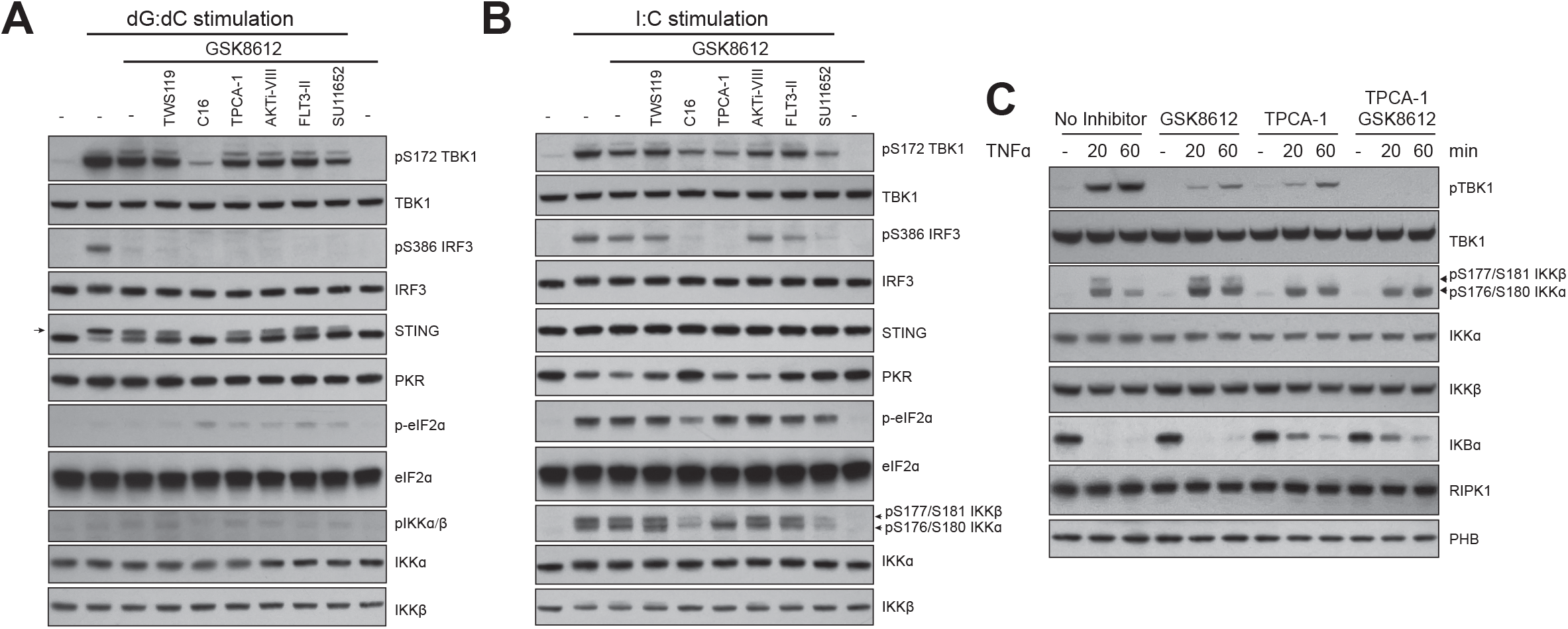
IKKꞵ contributes to the activation of TBK1 by dsRNA and TNFꭤ. **A** and **B**. Two-inhibitor experiments to explore upstream kinases involved in TBK1 phosphorylation. HT1080 cells were untreated, pretreated with GSK8612 (10µM) either alone, or GSK8612 plus one additional kinase inhibitors (TWS119, C16, TPCA-1, AKTi-VIII, FLT3-II and SU11652, 10µM for each condition) for 1hr. Cells were then stimulated with dsDNA (poly dG:dC, A) or dsRNA (poly I:C, B). Total cell lysates were prepared 2hrs later and the phosphorylation and expression of TBK1, IRF3, STING, PKR, eIF2ꭤ, IKKꭤ/ꞵ were determined by Western blotting with specific antibodies. Phospho-STING is labeled with an arrow in panel A. **C**. IKKꞵ cooperates with TBK1 for its activation after TNFꭤ stimulation. HT1080 cells were untreated or pretreated with TPCA-1 (10µM), GSK8612 (10µM) either alone or in combination for 1hr, followed by TNFꭤ (10ng/ml) stimulation for increasing time (20 to 60 minutes). Cell lysates were prepared after indicated time. The expression and phosphorylation profile of TBK1, IKKꭤ/ꞵ, IKBꭤ, RIPK1 and PHB were determined by Western blotting with specific antibodies.

Combined treatment of C16 and GSK8612 significantly inhibited IRF3 phosphorylation induced by dsRNA (Fig. 3B). TPCA-1, a specific inhibitor for IKKβ, also blocked dsRNA-induced IRF3 phosphorylation when used in combination with GSK8612 (Fig. 3B). Since PKR has been shown to be required for the activation of IKKα/β kinases (30, 31), and we observed that C16 strongly inhibited the dsRNA-induced phosphorylation of eIF2α - an established PKR target (Fig. 3B). These data suggest that the kinase responsible for TBK1-independent IRF3 phosphorylation induced by dsRNA is likely IKKβ. Monitoring phospho-IKKα/β profile in cells pretreated with various inhibitors subsequently stimulated with dsRNA revealed that TPCA-1 did block IKKβ phosphorylation (the disappearance of the upper band in the phospho-IKKα/β doublet, Fig. 3B). Interestingly, C16 and SU11652 inhibited both IKKα and β phosphorylation induced by dsRNA. As a result, C16, TPCA-1 and SU11652 significantly inhibited dsRNA-induced IRF3 phosphorylation when used in combination with GSK8612 (Fig. 3B). Although the impact of these inhibitors was not as dramatic as on IRF3 phosphorylation, they also reduced TBK1 phosphorylation when used in combination with GSK8612 followed by dsRNA stimulation (Fig. 3B). Thus, IKKβ also appears to contribute to dsRNA-induced TBK1 phosphorylation.

### IKKβ activity is required for TNFα-induced TBK1 phosphorylation

We also investigated the activation of TBK1 in response to TNFα signaling. A previous report showed that TBK1 activation by TNFα occurs solely through IKKα/β in mouse embryonic fibroblasts (MEFs) (21), as an inhibitor of IKKβ completely blocked TNFα-induced TBK1 phosphorylation in IKKα deficient MEFs (21). To determine whether the same is true of human cells, we treated HT1080 cells with TNFα for increasing time (20 and 60 minutes) in the absence/presence of TPCA-1 and GSK8612 either alone or in combination. Strikingly, these inhibitors alone reduced TBK1 phosphorylation, but their combination completely blocked TNFα-induced TBK1 phosphorylation (Fig. 3C). Like the experiments described in Fig. 3B, the effect of TPCA-1 on IKKβ appeared specific, as it abolished S177/S181 phosphorylation of IKKβ, but not IKKα phosphorylation (Fig. 3C). Although delayed, IKBα degradation was still induced by TNFα treatment in the presence of TPCA-1 (Fig. 3C). These results show that IKKβ is involved in TNFα-induced TBK1 activation. In contrast to MEFs, where TBK1 phosphorylation by IKKα/β is essential for its full activation, IKKβ, but not IKKα, cooperates with TBK1 for its full phosphorylation after TNFα stimulation in human cells.

### A 7DG-sensitive kinase(s) is involved in the activation of TBK1 in dsDNA-STING pathways

It is striking that C16 in combination with GSK8612 significantly inhibited dsDNA-induced TBK1 phosphorylation (Fig. 3A). C16 is known to directly inhibit the kinase activity of PKR, which we confirmed (Fig. 3B). We tested whether another small molecule PKR inhibitor, 7DG, can also inhibit TBK1 activation when used in combination with GSK8612. 7DG does not inhibit PKR kinase activity, instead, it binds PKR on its C-terminus and interferes with interactions between PKR and other proteins (32). Remarkably, our results show that 7DG is a more potent inhibitor when used in conjunction with GSK8612 and this combination strongly inhibits dsDNA and a STING activator-diABZI (33) induced TBK1 phosphorylation (Fig. 4A, Supplementary Fig. 3A). By contrast, C16 is not very effective in blocking diABZI-induced TBK1 and STING activation when used together with GSK8612 (Supplementary Fig. 3B). Unexpectedly, we found that C16 disrupted the structural integrity of Golgi, which likely contributes to its inhibition of dsDNA-induced TBK1 phosphorylation when used together with GSK8612 (Supplementary Fig. 3C), as Golgi is a central organelle in dsDNA-STING signaling.

**Figure 4.**
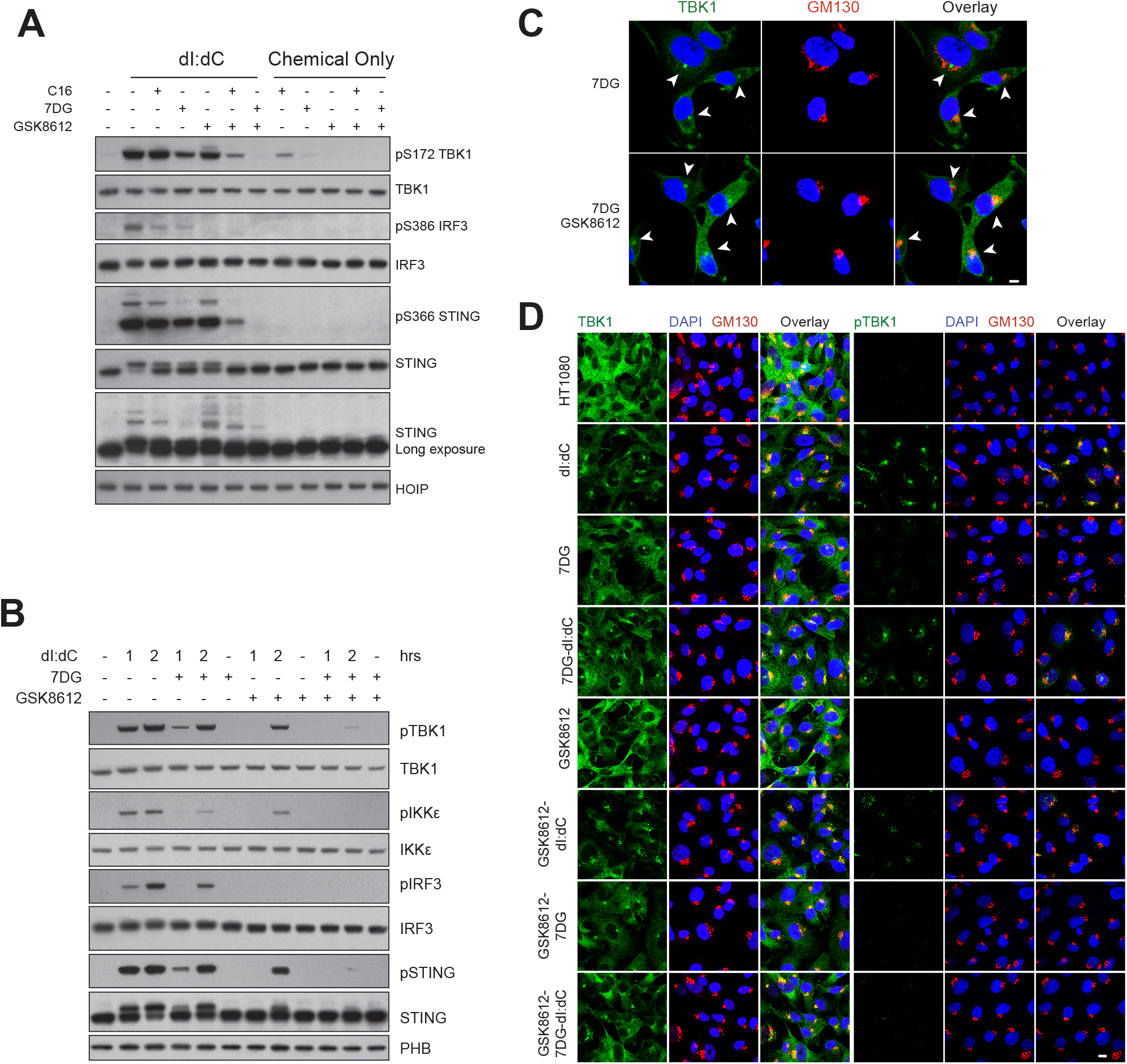
7DG in combination with GSK8612 potently inhibit dsDNA-induced TBK1 phosphorylation. **A**. 7DG is a more potent inhibitor of TBK1 activation when combined with GSK8612 compared to C16. HT1080 cells were untreated, pretreated with GSK8612, PKR inhibitor C16, 7DG (10µM for each inhibitor) either alone or together with GSK8612 and stimulated with dsDNA (poly dI:dC) for 2hrs. The expression and phosphorylation of TBK1, IRF3, STING and HOIP were determined by Western blotting with specific antibodies. **B**. 7DG in combination with GSK8612 inhibits phosphorylation of TBK1 induced by dsDNA. HT1080 cells were pretreated with 7DG or GSK8612 either alone or in combination for 1hr, and treated with dsDNA for increasing time (1 and 2hrs). Cell lysates were prepared after indicated time and the expression profile of various proteins were analyzed by Western blotting with specific antibodies. **C**. 7DG induces the perinuclear enrichment of a subset TBK1. HT1080 cells were treated with 7DG or 7DG/GSK8612 combination (10µM each) for 3hrs. Cells were fixed and stained for total TBK1 and GM130. Distributions of these proteins were analyzed by confocal microscopy. Scale bar: 10µm. **D**. HT1080 cells were pretreated with 7DG, GSK8612 either alone or in combination (10µM each) for 1 hr, and stimulated with dsDNA (poly dI:dC) for 2hrs. Cells were fixed and stained for total TBK1, pTBK1 together with GM130, and analyzed by confocal microscopy. Scale bar: 10µm.

We conducted kinetic studies of cells treated with dsDNA or diABZI for increasing time in the absence or presence of 7DG and GSK8612, either alone or in combination. For both dsDNA and diABZI, 7DG alone partially suppressed TBK1 phosphorylation, but its combined use with GSK8612, profoundly inhibited TBK1 and STING phosphorylation (Fig. 4B, Supplementary Fig. 3A). Since 7DG does not inhibit TBK1 kinase activity (Supplementary Fig. 4A), its inhibitory activity towards TBK1 activation must be through other kinases. Surprisingly, the effect of 7DG on TBK1 activation is PKR-independent, as knocking-down PKR expression by shRNA failed to change the cellular sensitivity to GSK8612 or 7DG (Supplementary Fig. 4B).

Next, we conducted IF experiments of cells treated with 7DG either alone or in combination with GSK8612 followed by dsDNA/diABZI stimulations, in order to gain insights into its inhibition of TBK1 phosphorylation. To our surprise, 7DG induced an unusual distribution of TBK1. As shown in Fig. 4C, staining cells treated with 7DG with an anti-TBK1 antibody revealed that a subset of TBK1 was enriched in a compact perinuclear organization, which was surrounded by Golgi apparatus (GM130 staining). The perinuclear enrichment of TBK1 induced by 7DG is not unique to HT1080 cells, as similar observations were made with multiple cell lines, including 293T and A549 cells (Supplementary Fig. 5). This peculiar TBK1 distribution may affect its association with signaling molecules on Golgi after dsDNA stimulation. 7DG also appears to affect the overall translocation of TBK1 after dsDNA stimulation, as the decline of TBK1 staining signal in other parts of the cytosol (away from Golgi) was less obvious in 7DG treated cells compared to control cells after dsDNA stimulation (Fig. 4D). GSK8612 in general does not affect the distribution of TBK1 induced by dsDNA or 7DG either alone or in combination (Fig. 4D).

**Figure 5.**
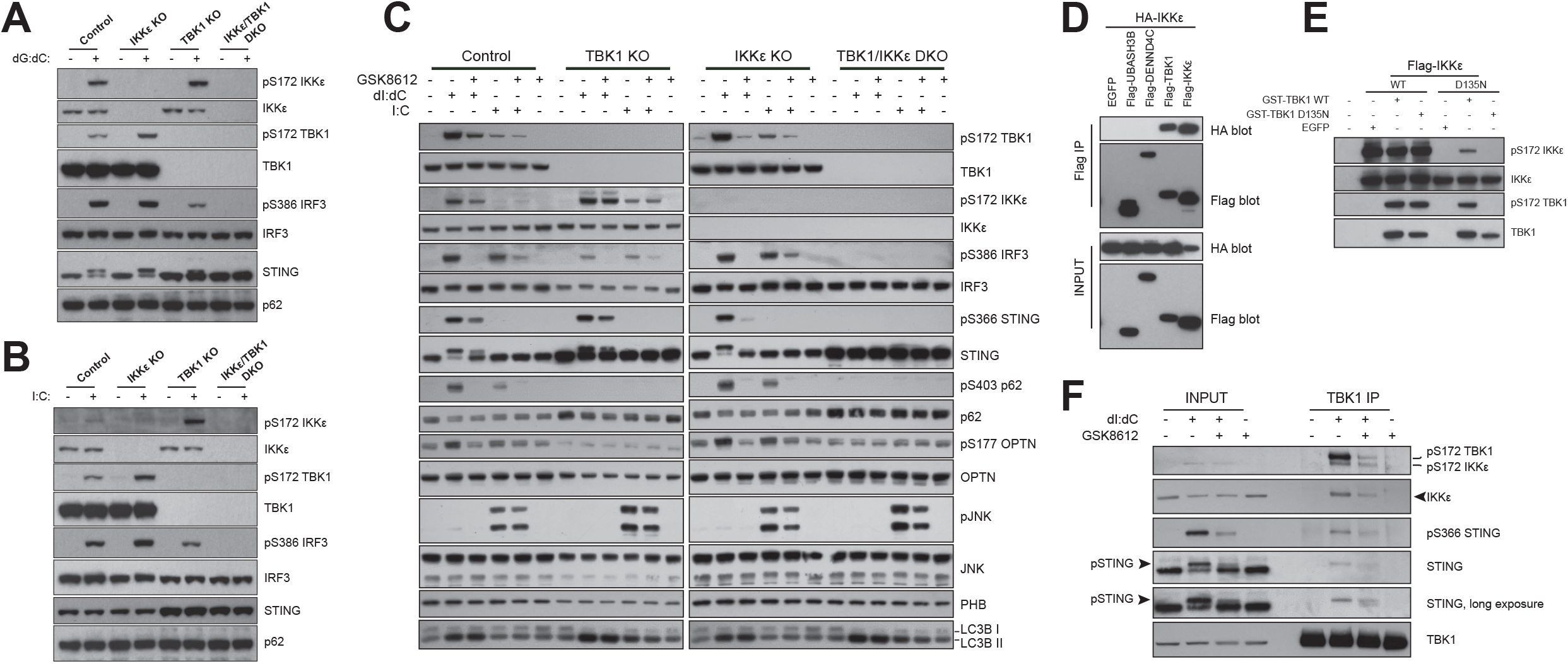
Competition and synergy between TBK1 and IKKε. **A**. and **B**. Contribution of TBK1 and IKKε to the activation of IRF3 in response to dsDNA and dsRNA. Control cells or TBK1 and IKKε single or double knockout cells were stimulated with dsDNA (A) or dsRNA (B) for 2hrs, total protein lysates were prepared, and the expression of various proteins and their phosphorylation profile were determined by Western blotting with specific antibodies. **C**. Control cells, TBK1 or IKKε single or double knockout cells were stimulated with dsDNA (poly dI:dC) or dsRNA (poly I:C) in the absence or presence of GSK8612 (10 µM, pretreated for 1hr) for 2hrs. Total protein lysates were prepared, and the expression and phosphorylation profile of various innate immune related proteins were determined by Western blotting with specific antibodies. **D**. Co-expressed TBK1 and IKKε interact with each other as revealed by immunoprecipitation experiments. Expression constructs for Flag-tagged UBASH3B, DENND4C, TBK1 and IKKε were co-transfected with a construct for HA-tagged IKKε into 293T cells. 24hrs later, cell lysates were prepared and subjected to immunoprecipitation with anti-Flag antibody (M2) conjugated beads. The associated HA-IKKε with the Flag-tagged proteins were analyzed by Western blotting with anti-HA antibodies. **E**. Co-expressed TBK1 and IKKε can phosphorylate each other. Expression constructs for wild type (WT) or kinase dead (D135N) IKKε were co-transfected with EGFP construct as a control or GST-tagged WT or kinase dead (D135N) TBK expression constructs. 24hrs later, total cell lysates were prepared, and the phosphorylation of S172 on TBK1 and IKKε determined by Western blotting with phospho-specific antibodies. **F**. dsDNA induces the interactions among TBK1, IKKε and STING. HT1080 cells were stimulated with dsDNA (poly dI:dC) in the absence or presence of GSK8612 (10µM, pretreated for 1hr) for 2hrs. Total protein lysates were prepared and subjected to anti-TBK1 antibody immunoprecipitation. The association of IKKε and STING with TBK1, and their phosphorylation status were determined by Western blotting with specific antibodies.

Despite the perinuclear enrichment of TBK1, pTBK1 staining signal in 7DG treated cells was barely detected (Fig. 4D). dsDNA-induced pTBK1 signal in 7DG treated cells was detected on the Golgi, and majority of this signal was due to autophosphorylation, as it was dramatically reduced by combined GSK8612 treatment (Fig. 4D). Remarkably, 7DG significantly reduced dsDNA-induced phosphorylation of TBK1 by upstream kinase(s) in cells treated with GSK8612 (to prevent TBK1 autophosphorylation) (Fig. 4D, compare the pTBK1 staining of GSK8612-dI:dC to GSK8612-7DG-dI:dC samples). These observations are in agreement with biochemical results of 7DG effect on TBK1 activation (Fig. 4A, 4B). We also observed that diABZI induced a highly Golgi-enriched TBK1 staining pattern in control cells (Supplementary Fig. 6). Neither 7DG nor GSK8612 prevent the enrichment of TBK1 on Golgi after diABZI treatment (Supplementary Fig. 6), but in combination they significantly inhibited TBK1 phosphorylation (Supplementary Fig. 6, panels in the far-right column). These results suggest that 7DG likely interferes with the initial S172 phosphorylation of TBK1 by upstream kinases, either by preventing the recruitment of the Golgi associated kinase to TBK1 or blocking its accessibility to TBK1.

**Figure 6.**
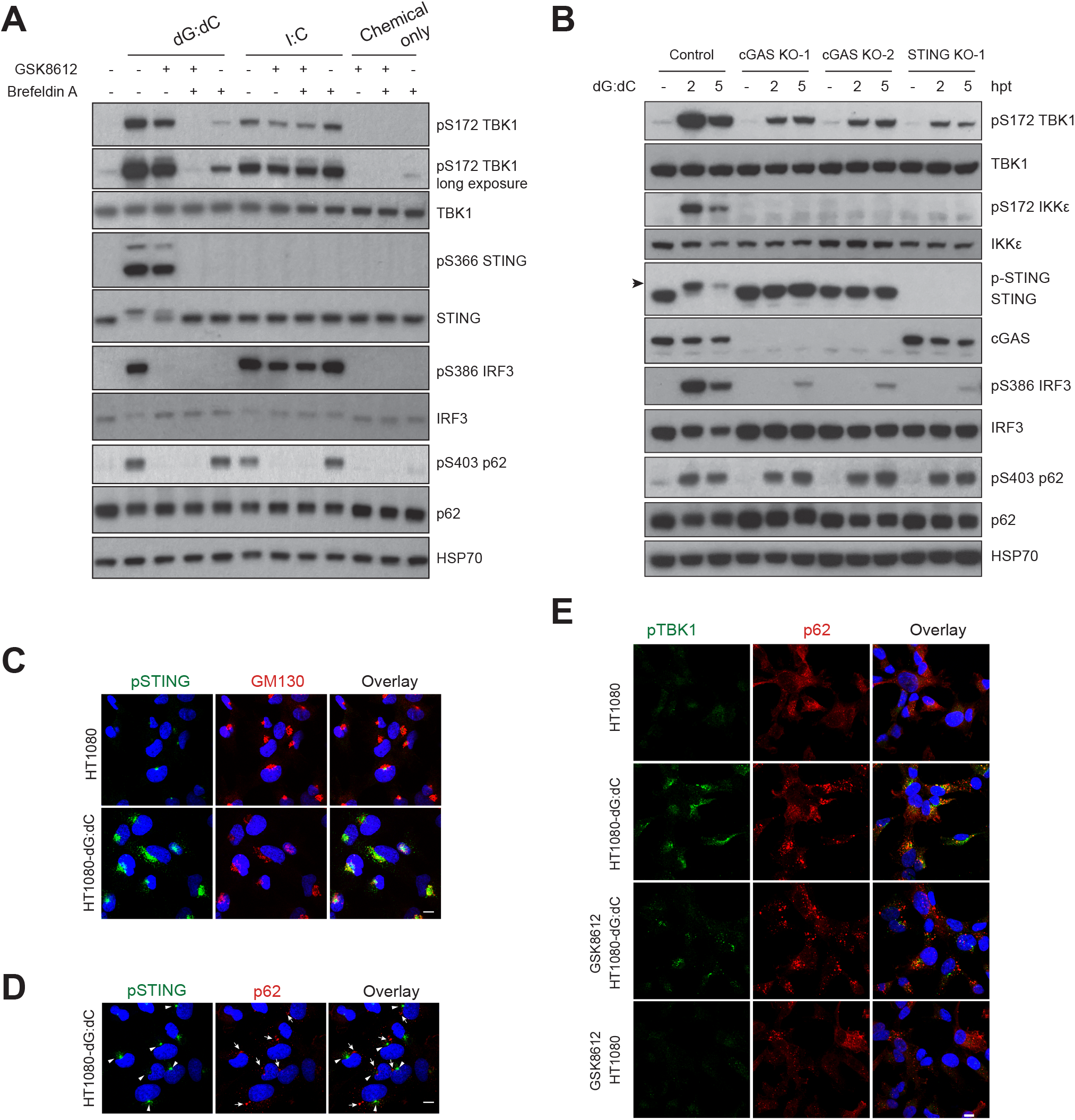
STING and p62 phosphorylation by TBK1 occurs at distinct signaling complexes. **A**. Brefeldin A uncouples TBK1-mediated STING/IRF3 phosphorylation from p62 phosphorylation. HT1080 were untreated, or pretreated with GSK8612 (10µM), BFA (5µM) either alone or in combination, then stimulated with dsDNA (poly dG:dC) or dsRNA (poly I:C). Total cell lysates were prepared 2hrs later and the expression and phosphorylation profile of various proteins were determined by Western blotting with specific antibodies. **B**. dsDNA-induced p62 phosphorylation is cGAS and STING independent. HT1080 cells of different genotype (control; cGAS KO, two lines; STING KO) were treated with dsDNA (poly dG:dC) for increasing time (2 and 5hrs). Total cell lysates were prepared and the expression and phosphorylation profile of TBK1, IKKε, STING, cGAS, IRF3 and p62 were determined by Western blotting with specific antibodies. **C**. S366 phosphorylated STING was enriched on Golgi after dsDNA stimulation. HT1080 cells were either untreated or stimulated with dsDNA for 2hrs. Cells were fixed and stained for phospho-S366 STING and GM130. Distributions of these proteins were analyzed by confocal microscopy. Scale bar: 10µm. **D**. Poor co-localization of pSTING with p62. HT1080 cells were stimulated with dsDNA for 2hrs. Cells were fixed and stained for pSTING and p62, and analyzed by confocal microscopy. Major signals of pSTING and p62 staining in some cells are labeled with arrowheads (pSTING) or arrows (for p62). Scale bar: 10µm. **E**. Partial co-localization of pTBK1 and p62. HT1080 cells were stimulated with dsDNA in the absence or presence of GSK8612 (10µM, pretreated for 1hr). 2hrs after stimulation, cells were fixed and stained for pTBK1 and p62. Their intracellular distributions were analyzed by confocal microscopy. Scale bar: 10µm.

### TBK1 and IKKε play distinct, partially overlapping roles in innate immunity

Our data clearly establish that upstream kinases are involved in TBK1 activation by both dsDNA and dsRNA. We note that, IKKε, a kinase closely related to TBK1, is also activated by dsDNA and dsRNA (S172 phosphorylation of IKKε, which is equivalent to TBK1 S172 phosphorylation, and is required for IKKε activation, Fig. 1D, 1E. dsDNA appeared to be a stronger inducer of TBK1 and IKKε activation compared to dsRNA in HT1080 cells). Genetic evidence suggests that TBK1 and IKKε play distinct, but sometimes redundant roles in cellular innate immunity (34-36). However, it is unclear whether TBK1 and IKKε are activated in a signal-specific manner and whether they cross-regulate each other. Also unknown is whether they display different sensitivities to inhibitors in vivo. To address these questions, we generated knockout cells for TBK1 and IKKε, either individually or in combination, by CRISPR-Cas9 technology, and challenged these knockout cells with dsDNA or dsRNA in the presence or absence of GSK8612.

We found that deleting either kinase gene alone did not lead to the absence of IRF3 phosphorylation after either dsDNA or dsRNA treatments (Fig. 5A, 5B). However, deletion of the *Tbk1* gene had a stronger effect on IRF3 phosphorylation, as the level of IRF3 S386 phosphorylation was considerably lower compared to the stimulated control or IKKε knockout cells (Fig. 5A-5C). We also noticed that the absence of one kinase led to a stronger activation of the other kinase compared to the level in control cells (Fig. 5A, 5B), indicating a competition between these kinases in dsDNA/RNA signaling pathways. Importantly, deleting both kinases in the double knockout cells abolished dsDNA/RNA-induced phosphorylation of IRF3 (Fig. 5A-5C). Moreover, p62 S403 and OPTN S177 phosphorylation occurred normally in IKKε knockout cells, but not in TBK1 knockout cells after dsDNA/RNA treatments (Fig. 5C). These data show that TBK1 and IKKε play partially redundant (for IRF3 phosphorylation) but unequal roles (IRF3, p62 and OPTN phosphorylation) in the activation of cellular innate immune response.

GSK8612 treatment revealed different outcomes in these knockout cells. In IKKε knockout cells where TBK1 activity is intact, GSK8612 not only abolished dsDNA-induced phosphorylation of IRF3, but also significantly reduced STING phosphorylation (Fig. 5C), revealing that TBK1 is strongly inhibited by GSK8612. dsRNA-induced IRF3 phosphorylation was similarly decreased by GSK8612 treatment compared to control cells (Fig. 5C). Our previous results showed that IKKβ is responsible for dsRNA-induced IRF3 phosphorylation when TBK1 is inhibited by GSK8612 (Fig. 3B). GSK8612 treatment of TBK1 knockout cells also inhibited dsDNA-induced IRF3 phosphorylation (Fig. 5C), which was carried out by IKKε and appeared weaker in TBK1 deficient cells (Fig. 5A, 5C). However, STING phosphorylation was less inhibited by GSK8612 in these cells compared to IKKε knockout cells (Fig. 5C). These results suggest that GSK8612 can also inhibit IKKε, but IKKε is less sensitive to this inhibitor compared to TBK1. This result is in agreement with published kinase selectivity data of GSK8612, as it targets TBK1 significantly better than IKKε from the chemoproteomics study (23).

### Cross-phosphorylation of TBK1 and IKKε

Comparing the level of pTBK1 in IKKε knockout cells pretreated with GSK8612 followed by dsDNA stimulation, to that in control cells revealed a significant decrease in IKKε knockout cells, indicating TBK1 trans-autophosphorylation accounts for a major activity in cells deficient for IKKε (Fig. 5C). This observation suggests that IKKε does contribute to TBK1 phosphorylation in dsDNA pathway, although it is not required for the activation of TBK1. On the other hand, treating control cells with GSK8612 followed by dsDNA/RNA stimulation did lead to a decreased level of pIKKε at S172 (Fig. 1D, 1E, 5C). Thus, TBK1 may also be involved in the phosphorylation of IKKε in these pathways. The combination of genetic tools and the TBK1 inhibitor revealed that TBK1 and IKKε not only compete, but also collaborate with each other in dsDNA/RNA pathways.

To gain further insights into the relationship between TBK1 and IKKε, we co-expressed these kinases in 293T cells. Immunoprecipitation (IP) of Flag-tagged TBK1 can pull down HA-tagged IKKε, demonstrating that these two kinases can exist in the same signaling complex (Fig. 5D). Co-expressed TBK1 and IKKε not only interact with each other, they are able to phosphorylate each other. As shown in Fig. 5E, co-expression of wild type (WT) TBK1 with a kinase dead (D135N) IKKε can lead to the specific S172 phosphorylation of IKKε, and vice versa. These results support the notion that TBK1 and IKKε may cross-activate each other in signaling events.

To demonstrate that TBK1 does, indeed, interact with IKKε in vivo, we conducted IP experiments with extracts from HT1080 cells that had been stimulated with dsDNA for 2hrs, a condition known to activate both TBK1 and IKKε (Fig. 1D, 5A, 5C). As shown in Fig. 5F, IP of total TBK1 protein can pull down IKKε in cells treated with dsDNA. Strikingly, this interaction is stimulation-dependent, as it was not detected in unstimulated cells. Interestingly, phosphorylated STING (the shifted band on SDS-PAGE, arrowhead), not the unphosphorylated form, can be detected in the IP sample from stimulated cells, suggesting that the TBK1-IKKε-STING complex is functional with components properly phosphorylated. Interestingly, TBK1 IP of extracts from cells pretreated with GSK8612 followed by dsDNA stimulation pulled down both phosphorylated and unphosphorylated STING (Fig. 5F), indicating TBK1-STING interaction is phosphorylation-independent. However, GSK8612 treatment reduced the amount of IKKε associated with TBK1, suggesting that TBK1-IKKε interaction is likely increased by phosphorylation.

Previous studies have shown that kinases can function in a structural role, independent of kinase activity in signaling events. For example, TNFα-induced NFκB signaling requires RIPK1 to engage with adaptor proteins downstream of TNFR1 (37). However, the kinase activity of RIPK1 is dispensable for this activity (38). We note that, although delayed and reduced, control and single knockout cells of either TBK1 or IKKε pretreated with TBK1 inhibitor still showed IRF3 phosphorylation upon dsRNA stimulation (Fig. 1E, 5C). This contrasts with TBK1 and IKKε double knockout cells, where dsRNA-induced IRF3 phosphorylation is abolished (Fig. 5A, 5C). These data show that the availability of the TBK1 or IKKε, irrespective of its enzymatic activity, could play a role in the recruiting of other kinases and engaging with downstream target proteins. The absence of TBK1 and IKKε prevents the assembly of signaling complexes capable of IRF3 phosphorylation. Our data thus provide an example in which TBK1/IKKε can also play a structural role in signal transduction, independent of kinase activity. This draws a distinction between kinase-dead missense mutation and TBK1 null mutation, and thus has implications in human diseases where missense loss-of-function and null mutations of TBK1 have been both reported, such as ALS/FTD (1).

### TBK1 engages target proteins in distinct subcellular complexes

Our data thus far reveal extensive crosstalk among TBK1, IKKε and IKKβ in various signaling pathways. Once activated, TBK1 engages and phosphorylates pathway-specific sets of target proteins. To gain insights into TBK1 target selectivity, we studied the phosphorylation of STING and p62 in the dsDNA pathway, as both are inducible and well-known TBK1 targets.

It is known that the translocation of STING from the ER to Golgi is required for the activation of the TBK1/IRF3 pathway after dsDNA stimulation (27, 39). Consistent with published results (27, 39), treating cells with brefeldin A (BFA), a drug that specifically blocks ER to Golgi transport, completely inhibited STING and IRF3 activation (Fig. 6A). All the activities on STING induced by dsDNA stimulation, such as phosphorylation and ubiquitination, were completely abolished by BFA treatment (Fig. 6A). As a result, IRF3 phosphorylation was also inhibited. Interestingly, BFA pretreatment followed by dsDNA stimulation did not completely prevent TBK1 phosphorylation, as a low level of pTBK1 was still induced by dsDNA. Surprisingly, pS403 p62 was still observed under these conditions, at a level similar to that observed with no BFA (Fig. 6A). BFA treatment thus effectively uncouples STING/IRF3 phosphorylation from p62 phosphorylation induced by dsDNA stimulation. When cells were pretreated with the combination of GSK8612 and BFA followed by dsDNA stimulation, IRF3, STING and p62 phosphorylation were all inhibited, demonstrating that target proteins of TBK1 have different sensitivities to chemical and subcellular perturbations (Fig. 6A).

In contrast to dsDNA, BFA had little effect on dsRNA-induced IRF3 phosphorylation. As TBK1, IRF3 and p62 phosphorylation were not affected at all by BFA treatment followed by dsRNA stimulation (Fig. 6A), demonstrating that dsRNA signaling does not involve ER to Golgi transport. Consistent with previous observations, although GSK8612 potently blocked dsRNA-induced p62 phosphorylation (Fig. 1A, 1E), it only reduced IRF3 phosphorylation irrespective of BFA treatment (Fig. 6A). These results clearly show that dsDNA and dsRNA use distinct signal transduction pathways and complexes for the activation of TBK1 and downstream signaling molecules.

To dissect the genetic components required for STING and p62 phosphorylation by TBK1, we generated knockout cells for cGAS and STING by CRISPR-Cas9 and challenged these cells by dsDNA transfection. As expected, deletion of cGAS abolished the phosphorylation of STING and its turnover and led to a significant reduction of pTBK1 after dsDNA stimulation. Strikingly, dsDNA-induced IKKε S172 phosphorylation was abolished by the deletion of either the *cGAS* or the *Sting* gene (Fig. 6B). However, TBK1 phosphorylation was still observed, despite the reduction, in the absence of either cGAS or STING. Surprisingly, dsDNA treatment led to a similar level of p62 phosphorylation in control or cGAS and STING knockout cells (Fig. 6B). These results show that although cGAS is important for dsDNA-induced activation of STING, it is dispensable for p62 phosphorylation by TBK1. Similarly, STING is not required for p62 phosphorylation induced by dsDNA stimulation. These data clearly demonstrate that dsDNA-induced phosphorylation of STING/IRF3 and p62 by TBK1 occur independently.

Physical separation of STING from p62 can be confirmed by IF staining. As shown in Fig. 6C, staining cells stimulated by dsDNA with an antibody recognizing S366-phosphorylated STING (pSTING) revealed its Golgi localization. This pSTING rarely co-localized with p62 despite the fact that both are phosphorylated by TBK1 (Fig. 6D). These results suggest that TBK1 is activated in distinct signaling complexes engaging with different target proteins in response to dsDNA stimulation. The observation that pTBK1 appears in a heterogeneous staining pattern also supports this conclusion (Fig. 2A, 2C)

Our data show that S403 of p62 is phosphorylated exclusively by TBK1 under dsDNA/RNA stimulations (Fig. 1A, 1D, 1E, 5C). To characterize features of this TBK1-specific target, we co-stained pTBK1 and p62. dsDNA induced a partial co-localization of pTBK1 with p62 (Fig. 6E) with some pTBK1 signals co-localizing with p62 signals in a punctate pattern. The majority of pTBK1 signal that did not co-stain with p62 was perinuclear, and likely on the Golgi apparatus based on our previous results (Fig. 2A, 2C). Interestingly, treating cells with GSK8612 did not prevent the partial co-localization of pTBK1 with p62 despite the reduction of pTBK1 signal (Fig. 6E), suggesting that TBK1 within these complexes was phosphorylated by other kinases as well. This observation is similar to the result that pTBK1 was observed on Golgi in cells treated with dsDNA in the presence of GSK8612 (Fig. 2C). Co-staining of pTBK1 with p62 in the presence of GSK8612 also confirms that TBK1 inhibition, rather than substrate accessibility, is responsible for the inhibition of p62 phosphorylation by GSK8612. We also showed that BFA treatment did not prevent the partial co-localization of pTBK1 with p62 after dsDNA treatment (Supplementary Fig. 7). Collectively, these results show that TBK1 is recruited to distinct signaling complexes after dsDNA stimulation, which largely determines its selective phosphorylation of specific target proteins. Importantly, the phosphorylation of TBK1 by upstream kinases in distinct signaling complexes appears to be a common step required for its activation.

**Figure 7.**
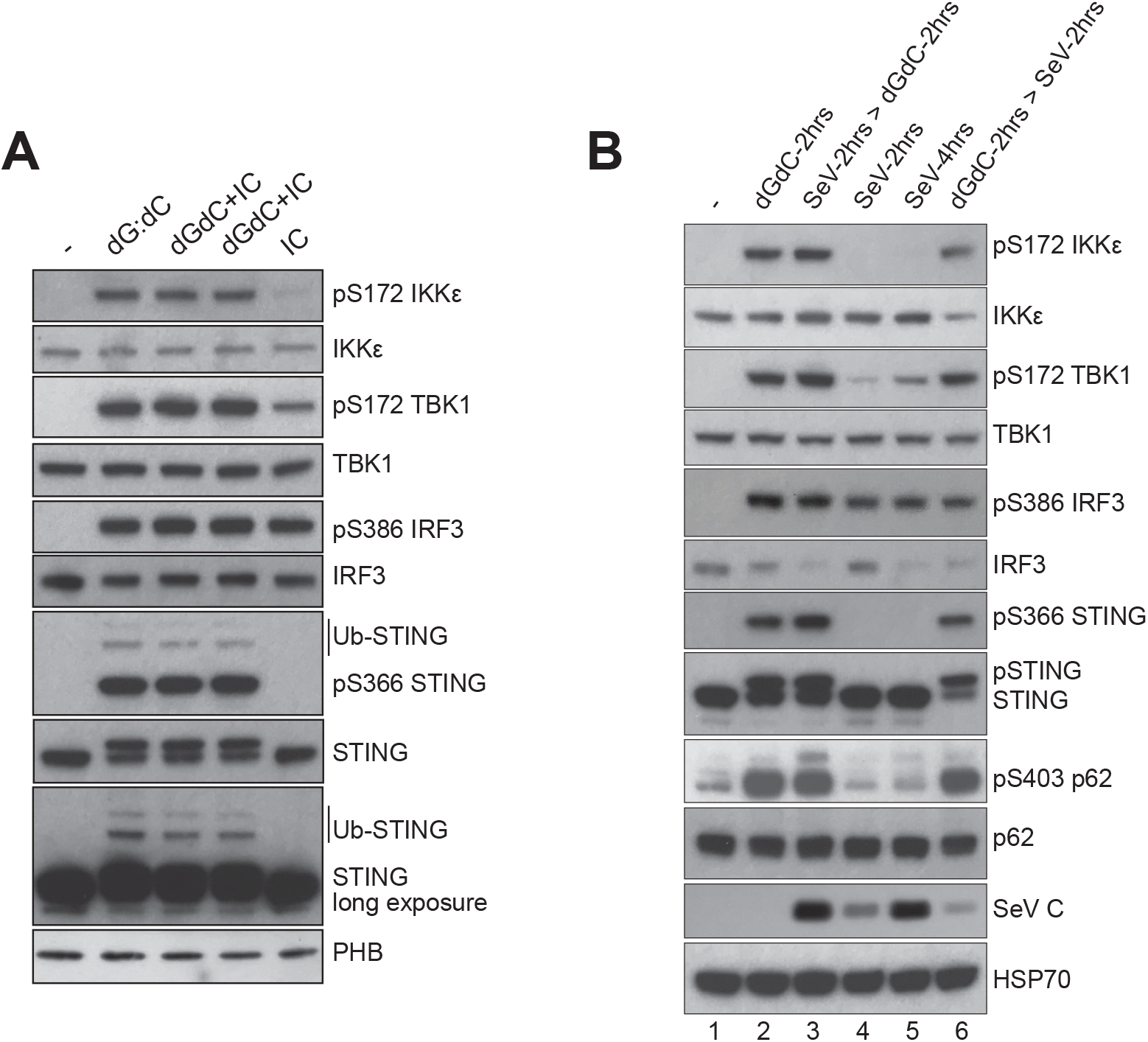
TBK1 activation by simultaneous or sequential stimulations. **A**. simultaneous stimulation of cells by dsDNA and dsRNA activates both pathways. HT1080 cells were treated with dsDNA (poly dG:dC), dsRNA (poly I:C) either alone or in combination for 2hrs. Total cell lysates were prepared and the expression of TBK1, IKKε, IRF3, STING and their phosphorylation profile were determined by Western blotting with specific antibodies. **B**. Activation of innate immunity by sequential infection and dsDNA stimulation. HT1080 cells were stimulated with dsDNA for 2hrs (lane 2), or infected with SeV for 2hrs followed by dsDNA treatment for another 2hrs (lane 3), infected with SeV for 2hrs (lane 4), 4hrs (lane 5), or stimulated by dsDNA for 2hrs followed by SeV infection for 2hrs (lane 6). Total cell lysates were prepared, and the expression and phosphorylation profile of selected proteins were determined by Western blotting with specific antibodies.

### Distinct TBK1 signaling complexes can be activated simultaneously or sequentially

Finally, we investigated the relationships between different TBK1 signaling complexes, e.g., whether the activation of one sequesters TBK1 and prevents the activation of the other. We conducted two types of experiments: cells were challenged with two different inducers either together, or sequentially. For the combined challenge, we stimulated cells with dsDNA and dsRNA together, and compared the levels of pTBK1, pIRF3 and pSTING to those from single treatment conditions. As shown in Fig. 7A, dsRNA (poly I:C) did not prevent the phosphorylation of STING induced by dsDNA. The combined stimulation of both inducers actually slightly increased the levels of pTBK1 and pIRF3 compared to single inducer treatment (Fig. 7A). For the sequential treatments, we challenged cells with dsDNA for 2hrs, then infected these cells with SeV for another 2hrs, or conducted infection 2hrs prior to dsDNA challenge. Similar to dsRNA treatment, SeV infection did not inhibit the activation of STING by dsDNA (Fig. 7B). Phosphorylation of p62 S403, which was barely induced by SeV, was robustly induced in cells challenged with dsDNA, even if they were infected with SeV 2hrs prior to dsDNA stimulation (Fig. 7B). These observations clearly show that TBK1 molecules within the same cell can be differentially activated in response to different stimuli, a feature that could be essential for cells to mount strong innate immune responses against combinations of pathogens.

## Discussion

Multiple studies have shown that TBK1 is activated by the phosphorylation of S172 within its kinase activation loop (10-12, 28, 40), and this appears to involve, at least in part, trans-autophosphorylation between distinct dimers (10, 11). However, as unphosphorylated TBK1 is inactive, there must be an upstream kinase or kinases that prime autophosphorylation in vivo. Here we show that TBK1 is activated by phosphorylation by an upstream kinase and by trans-autophosphorylation in cultured cells. The highly specific TBK1 inhibitor GSK8612, which potently inhibits TBK1 catalytic activity, does not prevent the phosphorylation of TBK1 S172 in cellular response to dsDNA, dsRNA, STING activator diABZI, TNFα stimulation, or SeV infection. Using dsDNA and dsRNA as examples, we show that these TBK1 activators induce a dynamic intracellular translocation of TBK1 within the cell, leading to its enrichment on Golgi and mitochondria, respectively. Significantly, GSK8612 does not inhibit TBK1 intracellular localization, nor the initial S172 phosphorylation in response to dsDNA/RNA, suggesting that TBK1 autophosphorylation is primed by upstream kinase(s), followed by amplification of kinase activity through trans-autophosphorylation.

### Involvement of distinct kinases in the phosphorylation of TBK1 in response to different inducers

Distinct upstream kinases are involved in TBK1 phosphorylation in response to different inducers (diagramed in Fig. 8). Utilizing genetic and chemical tools, we show that IKKβ contributes to dsRNA and TNFα-induced TBK1 activation, while IKKε partially contributes to dsDNA-induced TBK1 activation. Specifically, treating cells with the IKKβ inhibitor TPCA-1 in conjunction with GSK8612 abolishes TNFα-induced TBK1 S172 phosphorylation (Fig. 3C), and reduces pTBK1 induced by dsRNA treatment (Fig. 3B). However, the combined treatment of TPCA-1 and GSK8612 has a minimal effect on the induction of pTBK1 by dsDNA (Fig. 3A). By contrast, while GSK8612 in combination with C16 potently inhibits dsDNA-induced TBK1 phosphorylation (Fig. 3A), they only modestly reduce the level of pTBK1 induced by dsRNA (Fig. 3B). Thus, specific responses of TBK1 to distinct inducers appear to involve multiple upstream kinases (Fig. 8).

**Figure 8:**
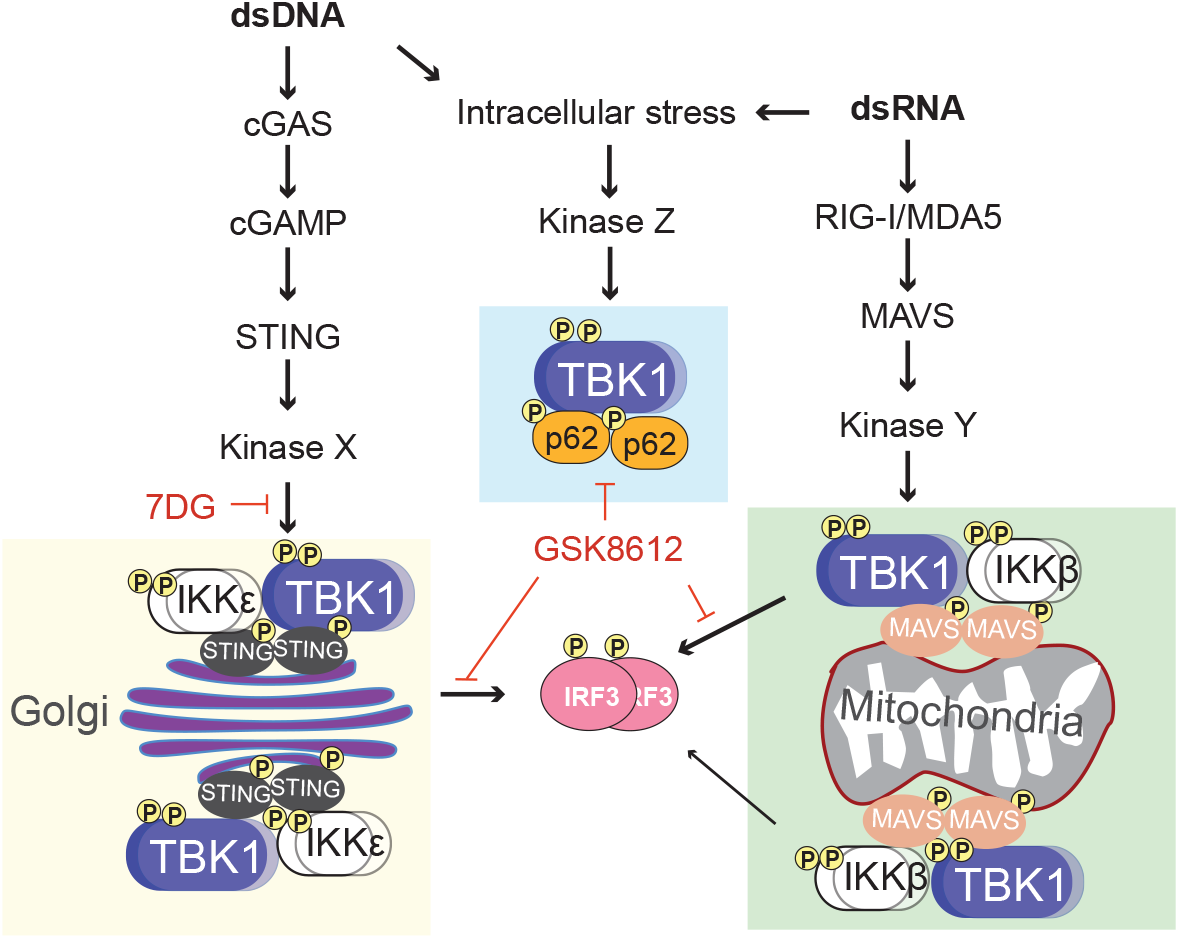
TBK1 activation by dsDNA and dsRNA in cultured human cells. Challenging cells with dsDNA induces cGAS-mediated production of the secondary messenger cGAMP, which binds to STING, leading to the translocation of STING from the ER to Golgi. TBK1 is activated along the way by a yet to be identified kinase(s) X, followed by TBK1 trans-autophosphorylation (not diagramed). Kinase X is sensitive to 7DG treatment. TBK1, STING and IKKε form signaling complexes on Golgi, and TBK1 and IKKε partially activate each other. Both activated TBK1 and IKKε can phosphorylate IRF3, and GSK8612 potently inhibits this phosphorylation. By contrast, intracellular dsRNA sensing by RIG-I/MDA5 activates MAVS on the mitochondria membrane. A yet to be identified upstream kinase(s) Y is involved in the initial phosphorylation of TBK1 followed by the amplification of the activation signal through trans-autophosphorylation (not diagramed). TBK1, MAVS and IKKꞵ form signaling complexes on mitochondria. IKKꞵ is partially involved in the activation of TBK1, it can also phosphorylate IRF3 (to a less extent compared to TBK1). GSK8612 inhibits TBK1-mediated IRF3 phosphorylation, but not IKKꞵ. Transfecting cells with dsDNA/RNA also induces cellular stress responses in the cytosol, which lead to the activation of TBK1 by an upstream kinase Z and the subsequent phosphorylation of p62 by TBK1. GSK8612 potently inhibits TBK1-mediated p62 phosphorylation.

Unlike kinase inhibitors used in this study, the small molecule 7DG does not inhibit TBK1 kinase activity. However, when combined with GSK8612, it significantly inhibited dsDNA and diABZI-induced TBK1 activation. It is striking that 7DG induces a perinuclear aggregation of TBK1 (Fig. 4D). Although this unique distribution of TBK1 does not appear to affect its trans-autophosphorylation, it likely hinders the recruitment of TBK1 to the STING signaling complex on Golgi. The lack of robust TBK1 phosphorylation in 7DG treated cells also showed that aggregation of TBK1 alone (as 7DG is capable of inducing TBK1) is not sufficient for TBK1 activation. 7DG experiments clearly demonstrate that both upstream kinases and TBK1 are required for full activation.

The data presented here also reveal crosstalk between TBK1 and its closely related kinase IKKε. We found that dsDNA potently activates both TBK1 and IKKε, and these two kinases not only compete, but also collaborate with each other in signaling (Fig. 5A-C, 5F). Through genetic and chemical manipulations, we showed that IKKε and TBK1 contribute to the phosphorylation of each other when induced by dsDNA and dsRNA. In contrast to canonical IKKs, where IKKα and IKKβ exist in a complex (41), TBK1 and IKKε are not recruited to the same complex prior to stimulation (Fig. 5F). We note that dsDNA induces the formation of signaling complexes containing TBK1, IKKε and STING, with STING fully phosphorylated. However, deleting only one kinase gene from the cell does not significantly impair STING phosphorylation, demonstrating a promiscuous and redundant relationship between these two kinases in STING signaling. In general, TBK1 is broadly involved in the phosphorylation of many cellular proteins in different signaling pathways, including innate immunity, autophagy and cell death. By contrast, IKKε appears to participate in selected pathways and may fine-tune the outcomes of those signaling events (Fig. 5A-5C).

The assembly of distinct signaling complexes in response to different inducers is an interesting feature of the mechanism of TBK1 activation, as TBK1 interacts with different upstream kinases and gains accessibility to unique sets of target proteins. Unexpectedly, we found that inhibiting TBK1 catalytic activity by GSK8612 does not necessarily lead to the inhibition of phosphorylation of its target proteins (such as IRF3). It appears that other kinases recruited to the signaling complex can potentially phosphorylate presumed TBK1 only substrates in the vicinity. TBK1 may thus play a structural role in the assembly of signaling complexes and facilitate access of target proteins by other kinases. Our studies thus provide an example in which contributions of the kinase activity and a structural role of TBK1 in signaling events can be uncoupled.

### TBK1 recruitment to distinct subcellular signaling complexes as a mechanism for substrate selectivity

We found that brefeldin A uncouples STING/IRF3 phosphorylation by TBK1 from TBK1-mediated p62 phosphorylation (Fig. 6A). Genetic and cellular imaging analysis showed that STING phosphorylation and p62 phosphorylation occur independently (Fig. 6B, 6D), which is consistent with the heterogeneous staining pattern of pTBK1 (Fig. 2A, 2C). These findings demonstrate that TBK1 is recruited to distinct subcellular signaling complexes after induction. It appears that the uptake of nucleic acids into the cell through cationic-lipids activates the TBK1-p62 complex, and dsDNA or dsRNA activates STING-TBK1 or MAVS-TBK1 signaling complexes on Golgi and mitochondria, respectively (Fig. 8). We found that treating cells with different inducers either simultaneously or sequentially leads to TBK1 activation and engagement with pathway-specific downstream targets (Fig. 7A, 7B). One stimulation does not appear to inhibit the activation of another pathway involving TBK1. While it remains to be determined whether the same observation can be made with broad signaling pathways, the examples presented here show that the TBK1 pool within the cell is capable of being activated by distinct inducers. Given the multiple roles of TBK1 plays in many different signaling pathways, it appears that subpopulations of TBK1 may respond to specific cellular cues.

TBK1 has recently been shown to play distinct pathophysiological roles in different cell types/tissues. For example, the loss of TBK1 in different tissues has different effects on disease progression in a mouse model of ALS (42). Given the finding that TBK1 is recruited to different signaling complexes after induction, it is likely that TBK1 mutations associated with human diseases contribute to disease pathology by influencing cellular physiological functions in a signaling complex-specific manner.

In summary, we show that TBK1 activation in cultured cells involves both upstream kinase priming and subsequent trans-autophosphorylation. Our study reveals an extensive crosstalk among TBK1, IKKβ, IKKε and other kinases in response to different inducers. We illustrate the signal-dependent and spatially regulated activation of TBK1, with the recruitment of TBK1 to specific signaling complexes also confers target protein selectivity.

## Materials and methods

### Cell culture, chemicals and reagents

HT1080, 293T, HeLa and A549 cells were purchased from ATCC. They were cultured in Dulbecco’s Modified Eagle Medium (DMEM) containing 10% Fetal Bovine Serum (FBS). GSK8612 was purchased from Axon Medchem; MRT67307, BX795, brefeldin A, C16, 7DG, TPCA-1, TWS119, AKTi-VIII, FTL3-II and SU11652 were purchased from Sigma. Poly dG:dC and poly dI:dC were from Sigma, and poly I:C was purchased from Invivogen. TNFα was purchased from Invivogen. diABZI was purchased from Selleck Chemicals. Recombinant TBK1 protein, expression constructs for TBK1 and IKKε were described before (28). Concentrated Sendai virus stock was purchased from Charles River Lab (Cantell strain) and used at a concentration of 300 HAU/ml for indicated infection time.

### Generation of TBK1, IKKε, cGAS and STING knockout cells

HT1080 cells were transfected with two modified px459 plasmids, each encodes CAS9 and a distinct gRNA sequence targeting the *Tbk1* gene (5’-gaagtgctctgcatcttggc-3’ and 5’-gctactgcaaatgtctttcg-3’), the *Ikbke* (IKKε) gene (5’-caccgttgcgggccttgtacacac-3’ and 5’-caccgagagcacagccaattacctg-3’, or utilizing a single plasmid for both CAS9 and two gRNAs for *cGAS* and *Sting* genes as described (43). Puromycin selection was initiated 24hrs after transfection and lasted for two days. About 20 single clones were picked and expanded from surviving cells and tested for the loss of specific gene product by Western blotting. The deletion of the alleles from the positive clones was further confirmed by PCR.

### Transfection, Immunofluorescence staining and confocal analysis

All transfection experiments were carried out with Lipofectamine 2000 transfection reagent (Invitrogen) according to manufacturer’s manual. For immunofluorescence staining, cells were grown on Poly D-lysine coated coverslips (Corning BioCoat) and treated under various conditions. Cells were then fixed in 4% formaldehyde (diluted by Phosphate-buffered saline (PBS) from the 16% stock solution) for 10 minutes and washed 3 times by PBS (10 minutes each), followed by permeabilization in 0.1% Triton X-100 (in PBS) for 10 minutes. After 3 times of PBS washing (10 minutes each), cells were blocked in a buffer of 1% BSA, 5% FBS in PBS for one hour, then stained in primary antibody solution at 4°C overnight. Cells were washed 3 times in PBS (20 minutes each) before incubating with Alexa Fluor-conjugated secondary antibodies for 1hr at room temperature. After extensive washes (3 times, 20 minutes each in PBS), coverslips were mounted with DAPI containing media, and subjected to confocal microscopy analysis with the Olympus FV1000 system.

### Western blots, Co-immunoprecipitation and antibodies

Total cell lysates were prepared by lysing washed cultured cells directly with the Laemmli sample buffer (2% SDS, 5% 2-mercaptoethanol, 0.0625M Tris-HCl, pH6.8, 10% glycerol and 0.002% bromophenol blue). Samples were heated at 95°C for 10 minutes before resolving on SDS-PAGE. After electrophoresis, proteins from the gel were transferred to a PVDF membrane (Bio-Rad), blocked with 5% milk in Tris-buffered saline with 0.1% Tween 20 (TBST), and incubated with various primary antibodies solutions at 4°C overnight. Washed membranes were incubated with HRP conjugated secondary antibody (Cytiva) at room temperature for 1hr. After extensive washes in TBST, specific protein bands on the membrane were visualized with ECL reagents (Thermo Scientific). For TBK1 Co-IP, cell lysates were prepared in a lysis buffer (20 mM Tris∙HCl, pH 7.5, 150 mM NaCl, 1% Triton X-100, 1 mM EDTA, 30 mM NaF, 1 mM glycerophosphate, 1X proteinase inhibitor (Roche), and 1 mM Na_3_VO_4_). Magnetic anti-Protein A beads (Sigma) were washed in a Beads Wash Buffer (BWB: 0.05% Tween 20 in PBS) for 3 times and incubated with anti-TBK1 antibody at room temperature for 1hr. Beads were then washed three times with the same BWB followed by one wash in the lysis buffer and incubated with extracted cell lysates at 4ºC overnight. Beads were collected with a DynaMag™-2 magnet (Invitrogen) and washed for 4 times with the same lysis buffer. Flag IP was similarly conducted, except that anti-Flag M2 magnetic beads (Sigma) were used. IP samples were denatured in the Laemmli sample buffer and subjected to SDS-PAGE for Western blot analysis. Anti-pS172 TBK1, TBK1, pS172 IKKε, IKKε, pIKKα/β, IKKα, IKKβ, A20, pS177 OPTN, OPTN, pS386 IRF3, IRF3, pS403 p62, pJNK, JNK, PKR, p-eIF2α, eIF2α, PARP1, PACT, cGAS, pS366 STING, and STING antibodies were purchased from Cell Signaling Technology; anti-p62 and LC3B antibodies were purchased from Abcam. Anti-Flag (M2) antibody was from Sigma, anti-HA and RNF31 (HOIP) antibodies were from Bethyl Laboratories. Anti-GFP, PHB and HSP70 antibodies were from Santa Cruz Biotechnology. Anti-GM130 was from BD Biosciences. Anti-TOMM20 antibody was from EMD Millipore. The anti-Sendai virus C protein antibody was a gift from Dr. Ganes Sen (Cleveland, USA).

### In vitro kinase assays

Kinase assays were conducted in a buffer system of 20mM HEPES, pH 7.5, 20mM β-Glycerophosphate, 0.1mM Na_3_VO_4_, 10mM MgCl_2_ and 50μM ATP, with recombinant TBK1 (2µg) and Flag-IRF3 (1µg) as a substrate at 30ºC for 1hr. Phosphorylation of IRF3 was analyzed by Western blotting probed with the anti-phospho-S386 IRF3 antibody.

### Lentivirus mediated shRNA knockdown

The pLKO.1 shRNA construct targeting PKR (5’-tcctggctcatctctttattc-3’) was obtained from Horizon Discovery. The preparation of lentivirus and the knockdown procedure were described before (44).

## Acknowledgements

We thank Erin Flaherty for comments on the manuscript, and Dr. Frank J. M. van Kuppeveld for CRISPR/Cas9 plasmids targeting cGAS and STING. This work was supported by a grant from Project ALS (PRALS 2017-09), and unrestricted funds.

## Notes

### Competing Interest Statement

The authors have declared no competing interest.

