## Supplementary figures for "Complex Intracellular Mechanisms of TBK1 Kinase Activation Revealed by a Specific Small Molecule Inhibitor"

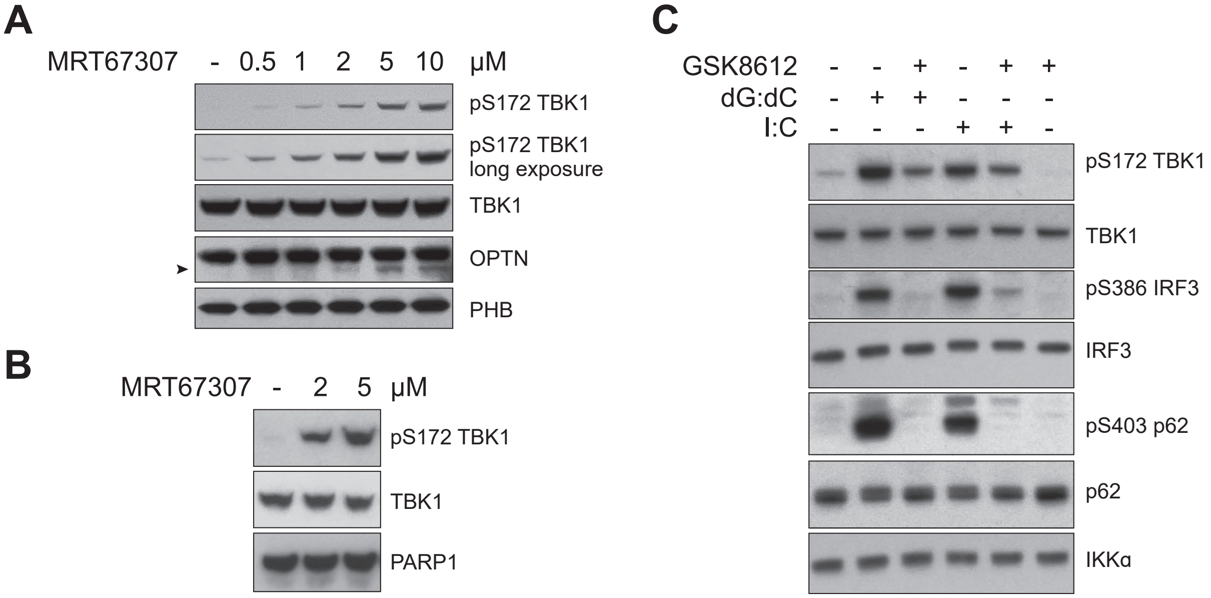


**Supplementary Figure 1. MRT67307 concentration-dependent induction of TBK1 S172 phosphorylation. A**. HT1080 cells were treated with increasing concentrations of MRT67307 (0.5μM to 10μM) for 3hrs, total cell lysates were prepared and the expression and phosphorylation of TBK1, OPTN, and PHB were analyzed by Western blotting with specific antibodies. A cleavage product of OPTN is labeled with an arrowhead. **B**. HeLa cells were treated with increasing concentrations of MRT67307 (2μM and 5μM) for 3hrs. Total cell lysates were prepared, and the expression of pTBK1, TBK1, PARP1 was analyzed by Western blotting with specific antibodies. **C**. HeLa cells were stimulated with dsDNA or dsRNA in the absence or presence of GSK8612 (cells pretreated with GSK8612 at 10μM for 1hr) for 2hrs. Total cell lysates were prepared and the expression and phosphorylation profile of TBK1, IRF3, p62 and IKKα were determined by Western blotting with specific antibodies.

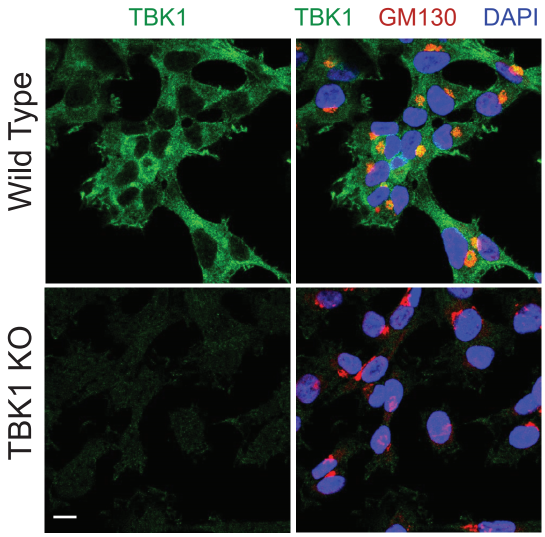


**Supplementary Figure 2. Validation of the anti-TBK1 antibody for immunofluorescent (IF) staining.** Wild type or TBK1 knockout (KO) HT1080 cells grown on coverslips were fixed and stained with anti-TBK1 antibody in conjunction with the anti-GM130 antibody at 4°C overnight. After standard IF procedures, samples were mounted with DAPI containing medium and subjected to confocal microscopy analysis. Scale bar: 10μm.

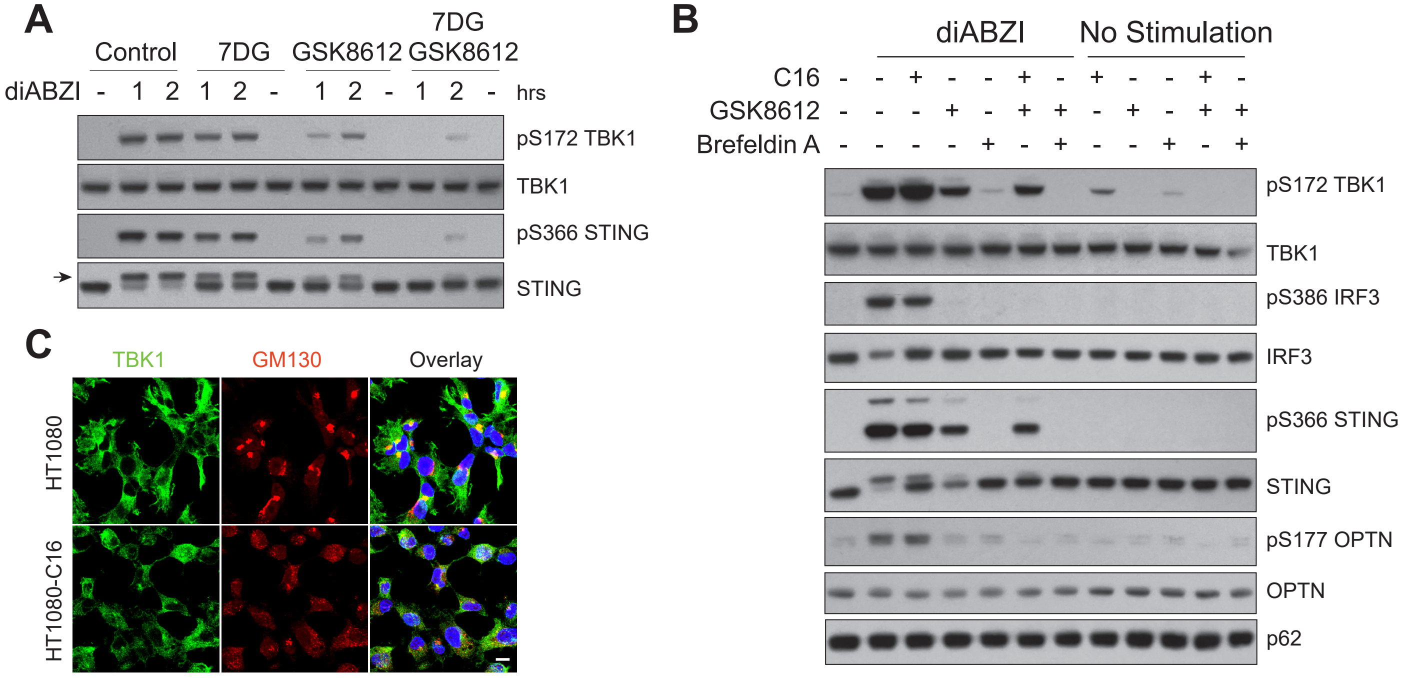


**Supplementary Figure 3. Effects of C16 and 7DG on TBK1 activation. A**. 7DG in combination with GSK8612 inhibits the phosphorylation of TBK1 induced by diABZI. HT1080 cells were pretreated with 7DG (10µM) or GSK8612 (10µM) either alone or in combination for 1hr, and treated with diABZI (10µM) for increasing time (1 and 2hrs). Cell lysates were prepared after indicated time and the expression and phosphorylation of TBK1 and STING analyzed by Western blotting with specific antibodies. Phospho-STING is indicated by an arrow. **B**. C16 is less efficient in inhibiting diABZI-induced TBK1 phosphorylation. HT1080 cells were untreated, pretreated with GSK8612 (10µM), PKR inhibitor C16 (10µM) and brefeldin A (BFA, 5µM) either alone or in different combinations, and stimulated with diABZI (10µM) for 2hrs. The expression and phosphorylation of TBK1, IRF3, STING, OPTN and p62 were determined by Western blotting with specific antibodies. **C**. C16 disrupts the structural integrity of the Golgi apparatus in HT1080 cells. HT1080 cells were either untreated or treated with C16 (10µM) for 3hrs. Cells were fixed and stained for TBK1 and GM130. Distributions of these proteins were analyzed by confocal microscopy.
Scale bar: 10µm.

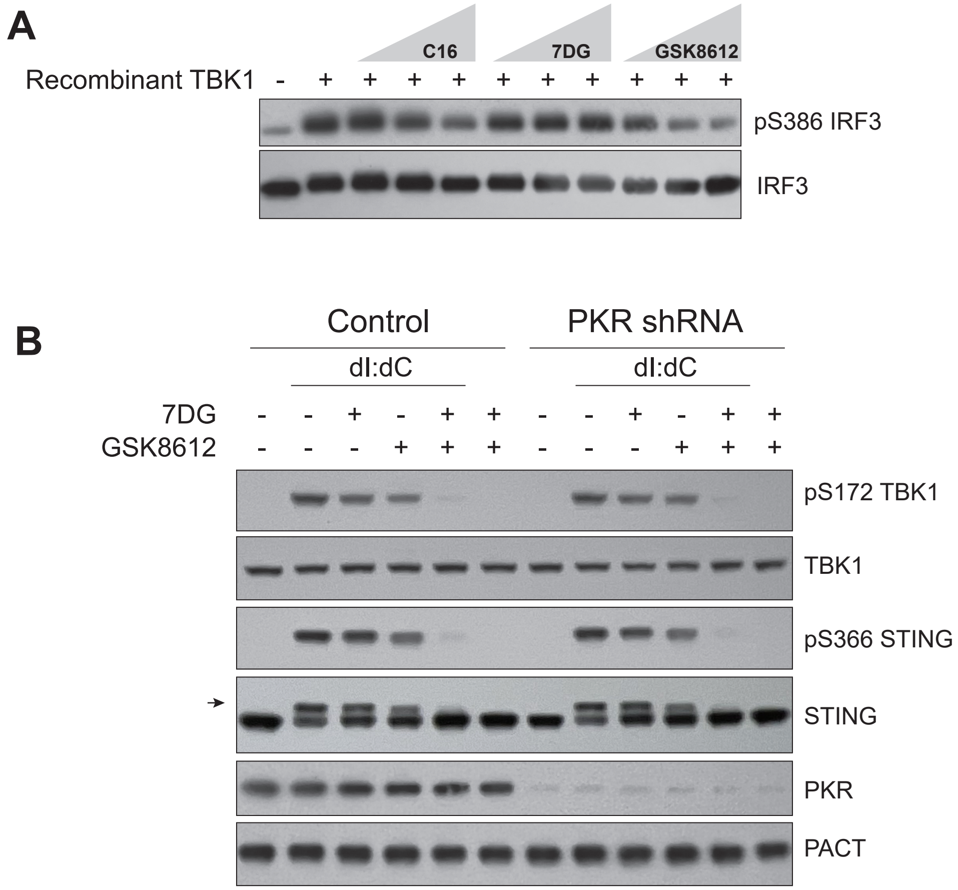


**Supplementary Figure 4. 7DG does not directly inhibit TBK1 kinase activity, and its effect on TBK1 phosphorylation is PKR-independent. A**. 7DG does not inhibit TBK1 kinase activity. Recombinant TBK1 protein was incubated with recombinant IRF3 in the absence or presence of increasing concentration of C16, 7DG and GSK8612 (1μM, 5μM and 20μM) at 30˚C for 1hr. The levels of phosphorylated and total IRF3 were determined by Western blotting with specific antibodies. **B**. Control or PKR knockdown HT1080 cells (by shRNA) were pretreated with 7DG (10μM), GSK8612 (10μM) either alone or in combination for 1hr, followed by dsDNA (poly dI:dC) treatment for additional 2hrs. Total cell lysates were prepared and the expression and phosphorylation profile of TBK1, STING, PKR and PACT were analyzed by Western blotting with specific antibodies. Phospho-Sting is indicated by an arrow.

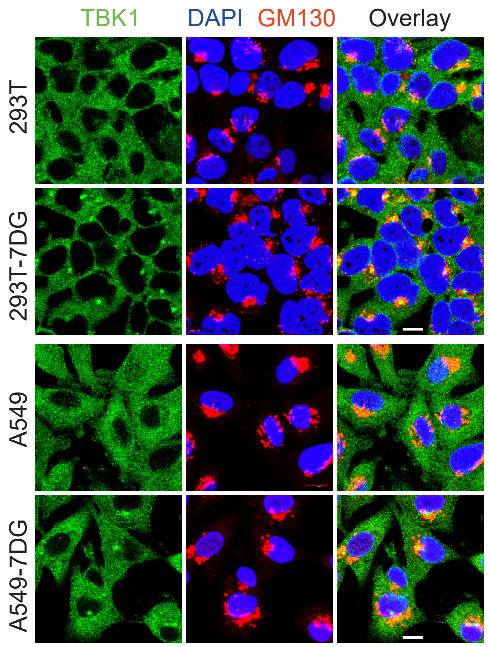


**Supplementary Figure 5. 7DG induces the perinuclear enrichment of TBK1 in multiple cell lines.** 293T and A549 cells grown on coverslips were treated with 7DG (10μM) for 3hrs, or left untreated. Cells were fixed and stained for TBK1 and GM130. The intracellular distribution of these proteins was analyzed by confocal microscopy. Scale bar: 10μm.

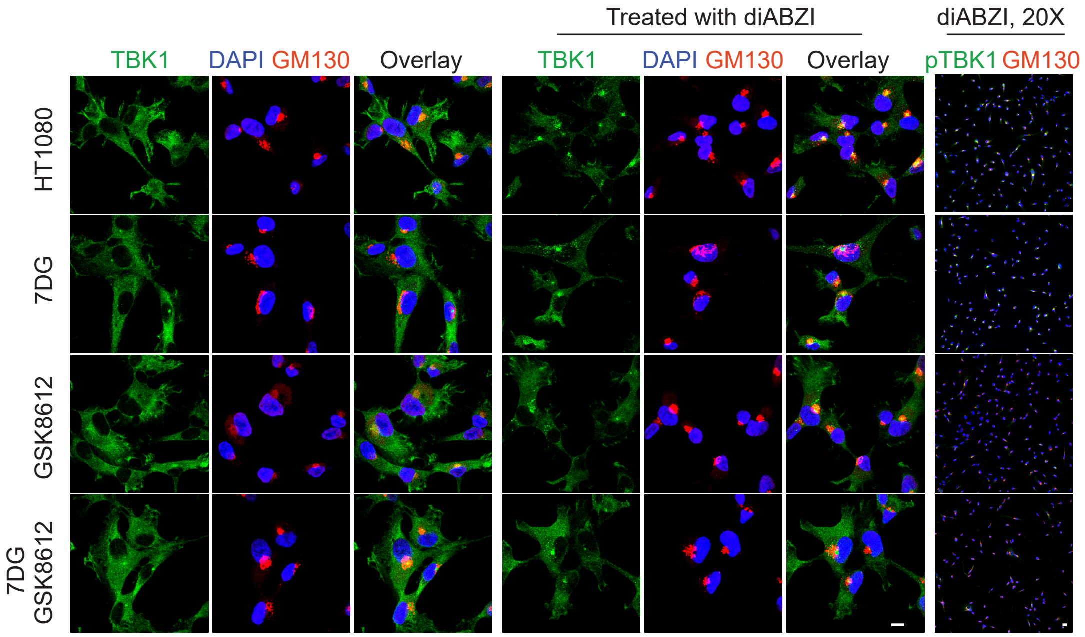


**Supplementary Figure 6. Effects of 7DG on the translocation of TBK1 induced by diABZI.** HT1080 cells were pretreated with 7DG (10μM), GSK8612 (10μM) either alone or in combination for 1hr, and stimulated with diABZI (10μM) for 2hrs. Cells were fixed and stained for total TBK1, pTBK1 together with GM130. Distributions of these proteins were analyzed by confocal microscopy. Scale bar: 20μm for the far-right panels, and 10μm for all other panels. The objective used to acquire the images in the far-right column (pTBK1 staining) was 20X, other images were acquired through a 60X objective.

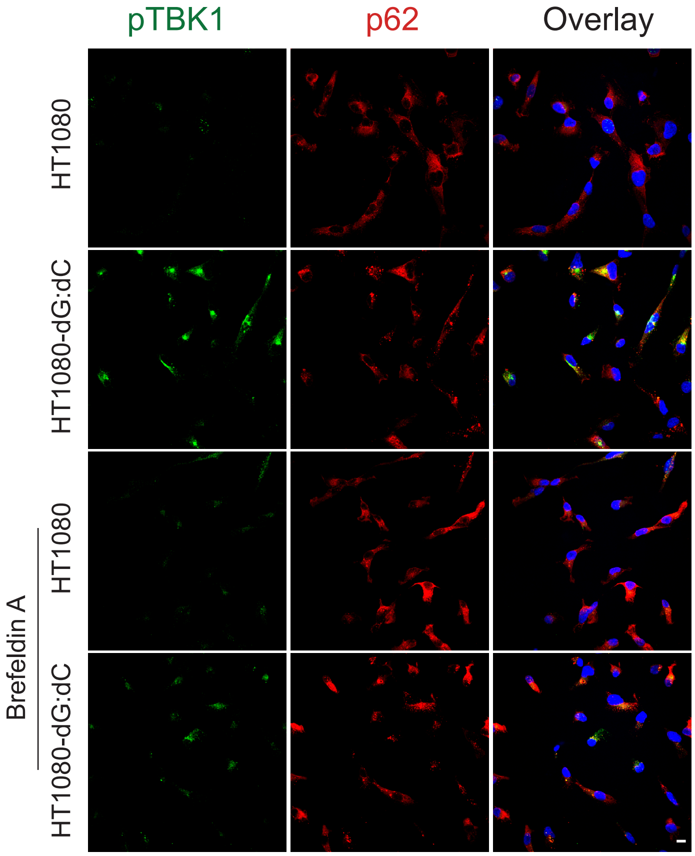


**Supplementary Figure 7. Partial co-localization of pTBK1 with p62 after dsDNA stimulation.** HT1080 cells were stimulated with dsDNA (poly dG:dC) for 2hrs in the absence or presence of BFA (5μM, pretreated for 1hr). Cells were then fixed and stained for pTBK1 and p62, and analyzed by confocal microscopy. Scale bar: 10μm.
